# Unraveling the link between Neuropathy Target Esterase NTE/SWS, lysosomal storage diseases, inflammation, and abnormal fatty acid metabolism

**DOI:** 10.1101/2023.08.11.552934

**Authors:** Mariana I. Tsap, Andriy S. Yatsenko, Jan Hegermann, Bibiana Beckmann, Dimitrios Tsikas, Halyna R. Shcherbata

## Abstract

Mutations in *Drosophila* Swiss Cheese (SWS) gene or its vertebrate orthologue Neuropathy Target Esterase (NTE) lead to progressive neuronal degeneration in flies and humans. Despite its enzymatic function as a phospholipase is well-established, the molecular mechanism responsible for maintaining nervous system integrity remains unclear. In this study, we found that SWS is present in surface glia that form the blood-brain-barrier (BBB) and that SWS is important to maintain its structure and permeability. Importantly, BBB glia-specific expression of *Drosophila* SWS or human NTE in the *sws* mutant background fully rescues surface glial organization and partially restores BBB integrity, suggesting a conserved function of NTE/SWS. Interestingly, *sws* mutant glia showed abnormal organization of plasma membrane domains and tight junction rafts accompanied by the accumulation of lipid droplets, lysosomes, and multilamellar bodies. Since the observed cellular phenotypes closely resemble the characteristics described in a group of metabolic disorders known as lysosomal storage diseases (LSDs), our data established a novel connection between NTE/SWS and these conditions. We found that mutants with defective BBB exhibit elevated levels of fatty acids, which are precursors of eicosanoids and are involved in the inflammatory response. Also, as a consequence of a permeable BBB, several innate immunity factors are upregulated. Treatment with anti-inflammatory agents prevents the abnormal architecture of the BBB, suggesting that inflammation contributes to the maintenance of a healthy brain barrier. Since a defective BBB is associated with many neurodegenerative diseases, a better understanding of the molecular mechanisms of inflammation may help to promote the use of anti-inflammatory therapy for age-related neurodegeneration.

## INTRODUCTION

Aging is the major risk factor for neurodegenerative disorders, which are a group of disorders characterized by the progressive degeneration and dysfunction of the nervous system, which includes Alzheimer’s disease, Parkinson’s disease, amyotrophic lateral sclerosis, frontotemporal dementia, and many others. These diseases typically result in the gradual loss of cognitive function, movement control, and other neurological functions. The exact causes of neurodegenerative diseases are often complex and not fully understood, but they can involve a combination of genetic, environmental, and lifestyle factors. There is a growing appreciation that inflammation plays an important role in age-related neurodegenerative diseases. Moreover, inflammation results in oxidative stress which contributes to pathogenesis of the age-related neurodegenerative diseases (Liu et al., 2018; Liu et al., 2017; Rojas-Gutierrez et al., 2017; Zuo et al., 2019). Older organisms frequently develop a chronic inflammatory status, a condition often named inflammaging (Chitnis and Weiner, 2017; Franceschi et al., 2000; Franceschi et al., 2018; McGeer and McGeer, 2004). One feature associated with neuroinflammatory degenerative diseases is dysfunction of the blood-brain barrier (BBB) (Takata et al., 2021). BBB disruption is detected in patients with such disorders as Alzheimer’s and Parkinson’s disease, amyotrophic lateral sclerosis and others (Blyth et al., 2009; Gray and Woulfe, 2015; Munji et al., 2019; Spencer et al., 2018; Sweeney et al., 2018; Whitson et al., 2022; Zhou et al., 2023). Moreover, many neurodegenerative diseases such as Alzheimer’s and Parkinson’s disease have been linked to dysfunction of lysosomal pathways (Issa et al., 2018). The lysosome-endosomal system is tightly associated with the maintenance of cell homeostasis and viability, regulation of cell death, oncogenesis, autophagy and inflammation (Peng et al., 2019). Therefore, there is potential for the use of anti-inflammatory therapy to treat neurodegenerative disorders. However, the molecular mechanisms underlying inflammaging remain unclear.

Human age-related neurodegenerative diseases can be accelerated by different stresses, which include a wide array of factors such as infection, trauma, diet, or exposure to toxic substances. Interestingly, abnormalities in the human Neuropathy Target Esterase (NTE), encoded by PNPLA6 (Patatin Like Phospholipase Domain Containing 6) gene are linked to both neurodegeneration types: toxin-induced and hereditary. NTE is a transmembrane protein anchored to the cytoplasmic face of the endoplasmic reticulum and acts as a phospholipase that regulates lipid membrane homeostasis (Glynn, 2005; Lush et al., 1998; Read et al., 2009).

Continuous inhibition of NTE activity by the organophosphorus compound tri-ortho-cresyl phosphate (TOCP) causes axonal degeneration in the central nervous system (CNS) and peripheral nervous system (PNS), a neuropathy that was consequently named Organophosphate-Induced Delayed Neuropathy (OPIDN) (Richardson et al., 2020; Richardson et al., 2013). Moreover, mutations in the NTE gene cause Gordon-Holmes or Boucher-Neuhäuser syndromes (Deik et al., 2014; Synofzik et al., 2014; Synofzik et al., 2015; Topaloglu et al., 2014) and a motor neuron disease called hereditary spastic paraplegia type 39 (HSP 39), in which distal parts of long spinal axons degenerate, leading to limb weakness and paralysis (McFerrin et al., 2017; Rainier et al., 2008). Genetically, HSP classification is based on the genes of origin called Spastic Paraplegia Genes, which is a large group (>80) of genes (Fereshtehnejad et al., 2023). Over the past few years, research has shown that HSP is associated with endo-lysosomal system abnormalities (Allison et al., 2017; Chang et al., 2014; Lim et al., 2015; Namekawa et al., 2007; Renvoise et al., 2014).

For various obvious reasons, humans are not ideal subjects for age-related research. Modelling human neurodegenerative diseases in various model organisms can provide us with needed knowledge about the first hallmarks of neurodegeneration and also signaling mechanisms that are disrupted upon aging. It was shown that NTE is widely expressed in the mouse brain, and its activity is essential for lipid homeostasis in the nervous system (Glynn et al., 1998; Moser et al., 2000). NTE deficiency results in the distal degeneration of the longest spinal axons, accompanied by swelling that encompasses accumulated axoplasmic material (Read et al., 2009). Specific deletion of NTE in the neuronal tissue induces neurodegeneration (Akassoglou et al., 2004). Despite its known molecular function, the mechanism by which it maintains nervous system integrity during hereditary and toxin-induced neurodegeneration remains unknown. *Drosophila melanogaster* is an excellent genetic model organism to investigate the molecular mechanisms of age-dependent neurodegenerative diseases, and it has been widely used to identify potential drug targets against neurodegenerative diseases (Jafari, 2010; Khan et al., 2019; Maitra and Ciesla, 2019; Maitra et al., 2019; Piper and Partridge, 2018; Prussing et al., 2013). Moreover, the fly nervous system is a great system to shed light on the evolutionarily conserved signaling pathways underlying disease pathology. In *Drosophila*, more than 70% of genes related to human diseased are conserved (Ugur et al., 2016). Studying human disease-related genes in *Drosophila* avoids the ethical issues of biomedical research involving human subjects. Together, *Drosophila* satisfies many of the requirements to study human diseases that allows not only dissection on cellular and molecular levels but also investigation of behavior and neurodegeneration during aging (Carney et al., 2023; Contreras and Klambt, 2023; Hajji et al., 2019; Ma et al., 2022; Rahmani et al., 2022; Yatsenko et al., 2021; Yatsenko and Shcherbata, 2021; Yon et al., 2020).

In this study, we used a *Drosophila* NTE/SWS model for human neurodegeneration. Swiss Cheese (SWS) protein is a highly conserved lysophospholipase that can regulate phosphatidylcholine metabolism (Lush et al., 1998; Zaccheo et al., 2004). It was also shown that SWS can act as a regulator of the PKA-C3 catalytic subunit of protein kinase A (Bettencourt da Cruz et al., 2008; Wentzell et al., 2014). NTE/SWS has been shown to result in lipid droplet accumulation, which is involved in neurodegeneration pathogenesis (Chang et al., 2019; Farmer et al., 2020; Melentev et al., 2021). Loss of *sws* leads to age-dependent neurodegeneration (Fig. 1B, arrows), CNS vacuolization and abnormal glial morphology accompanied by the formation of multilayered glial structures in the adult brain (Dutta et al., 2016; Kretzschmar et al., 1997). Recent studies have shown that the pan-glial knockdown of *sws* leads to increased levels of reactive oxygen species (ROS), which in turn induces oxidative stress (Ryabova et al., 2021). However, the role of SWS in distinct glial types is not clearly understood.

**Figure 1.**
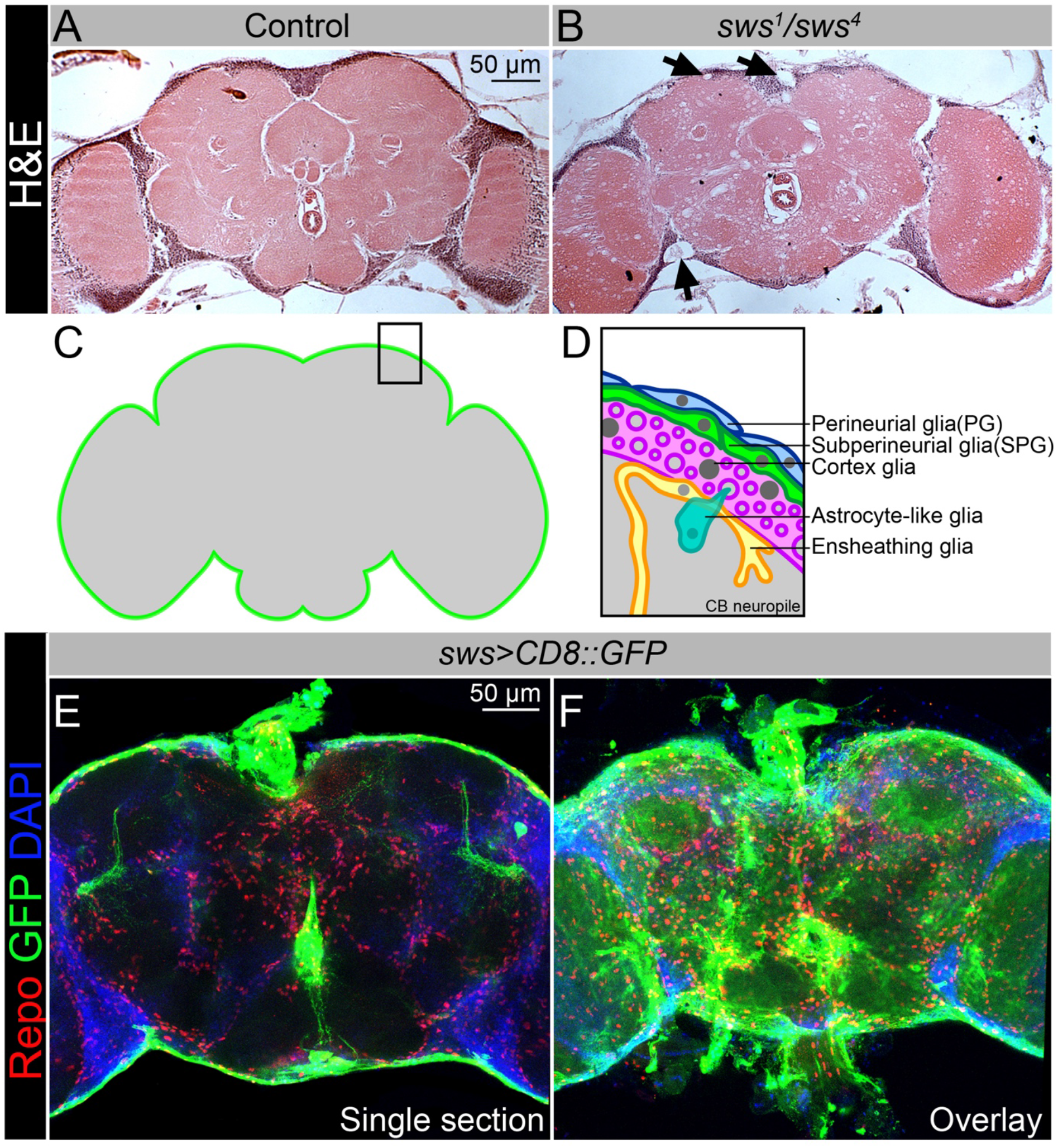
SWS expression pattern in *Drosophila* brain. **A-B.** Hematoxylin and eosin (H&E)-stained paraffin-embedded brain sections of the 30-day-old control *Oregon R x white^1118^* (**A**) and 30-day-old *sws^1^/sws^4^* transheterozygous flies (**B**). Arrows indicate neurodegeneration at the brain surface. Scale bar: 50 µm. **C-D.** Schemes of glia organization in the adult *Drosophila* brain - Perineurial glia (PG, blue), Subperineurial glia (SPG, light green), Cortex glia (pink), Astrocyte-like glia (turquoise) and Ensheathing glia (yellow). **E-F.** Expression pattern of *sws-Gal4* determined by combining of the transcriptional activator Gal4 under control of the *sws* gene promotor (*sws-Gal4*) and the *UAS-CD8::GFP* construct. Fluorescence images of the brain show that *sws* is expressed in all brain cells and strongly expressed in the surface glia (**E** - single section, **F** - Z-stack maximum projection). For SWS antibody staining pattern see Suppl. Figure 1. Glia cells are marked with Repo (red), *sws* expression is marked by the membrane CD8::GFP (green), and nuclei are marked with DAPI (blue). Scale bar: 50 µm.

Similar to multiple other organisms, the *Drosophila* nervous system is composed of neurons and glial cells. Commonly recognized nomenclature identifies six distinct glial cell types based on morphology and function: perineurial and subperineurial glia, cortex glia, astrocyte-like and ensheathing glia, and finally the PNS-specific wrapping glial cells (Trebuchet et al., 2019; Yildirim et al., 2019). All organisms with a complex nervous system developed BBB to isolate their neurons from blood (Limmer et al., 2014). In higher vertebrates, this diffusion barrier is established by polarized endothelial cells that form extensive tight junctions (Armulik et al., 2010), whereas in lower vertebrates and invertebrates the blood-brain barrier is entirely formed by glial cells, which are additionally sealed by septate junctions (Limmer et al., 2014). The *Drosophila* BBB includes two glial cell layers: the perineurial glial cells (PG) are primarily involved in nutrient uptake, whereas the main diffusion barrier is made by the subperineurial glia (SPG), which form pleated septate junctions (Babatz et al., 2018; Kremer et al., 2017; Schwabe et al., 2017).

Here we showed that NTE/SWS is present in the surface glia of *Drosophila* brain that form the BBB and that NTE/SWS is important for the integrity and permeability of the barrier. Importantly, glia-specific expression of *Drosophila* SWS or human NTE in the *sws* mutant background fully rescues surface glial organization and partially restores BBB integrity, suggesting a conserved function of NTE/SWS. An important observation upon *sws* deficit was the formation of intracellular accumulations within lysosomes, which is a characteristic feature of lysosomal storage disorders. Additionally, SWS regulates lipid metabolism, distribution of cell junction proteins, and organization of membrane rafts in BBB glia. Moreover, our research revealed that mutants with defective BBB exhibit elevated levels of several innate immunity factors as well as free fatty acids, which are known to play a role in inflammatory pathways. Importantly, the BBB phenotype can be alleviated by the administration of anti-inflammatory agents. These findings emphasize the complex interplay between SWS, BBB function, inflammation, and innate immunity, providing potential avenues for therapeutic interventions in related disorders.

## RESULTS

### SWS is expressed in the surface glia of Drosophila brain

Our previous data showed that NTE/SWS function is important for both glia and neuronal cells in the brain (Melentev et al., 2021; Ryabova et al., 2021). After downregulation of NTE/SWS in neurons, adult flies show a decrease in longevity, locomotor and memory deficits, and severe progression of neurodegeneration in the brain (Melentev et al., 2021). We have shown that NTE/SWS plays a role in the development of the learning center of the brain involved in short-term and long-term memory storage, olfactory control, and startle-induced locomotion (Melentev et al., 2021). In addition, we found that flies with NTE/SWS deficiency in neurons or glia show mitochondrial abnormalities as well as accumulation of ROS and lipid droplets (Melentev et al., 2021; Ryabova et al., 2021). Now we have decided to determine the cell type in which SWS plays a determining role in the maintenance of brain health.

Similar to its human counterpart, NTE, which is found in virtually all tissues, including the nervous system (https://www.proteinatlas.org/ENSG00000032444-PNPLA6/tissue), SWS is ubiquitously expressed in *Drosophila* brain, detected by immunohistochemical analysis using SWS-specific antibodies (Suppl. Fig. 1). SWS is a transmembrane phospholipase anchored to the cytoplasmic side of the endoplasmic reticulum to regulate lipid membrane homeostasis. Its cytoplasmic localization makes it difficult to determine precisely in which brain cell type it has more pronounced expression, as neurons and glia have very complex shapes and forms. Therefore, additional markers must be used to discriminate SWS expression in the brain. To address this, we overexpressed membrane-bound GFP under control of the *sws* promoter (*sws-Gal4; UAS-CD8::GFP*), which allows labeling of membranes of cells in which the *sws* promoter is active. Importantly, *sws* was strongly expressed in the glia that surround the brain and form the blood-brain selective permeability barrier (Fig. 1E, F). In *Drosophila*, the BBB is entirely made by two glial cell layers: perineurial glia and subperineurial glia (PG and SPG, Fig. 1C-D). With the help of sophisticated septate junctions, SPG cells form a tight barrier that prevents paracellular diffusion and separates the central nervous system from hemolymph. Since the BBB is protecting the brain from toxic substances, and SWS/NTE deregulation is associated with toxicity-induced neurodegeneration, we investigated whether SWS has a functional role in BBB maintenance and its selective permeability.

### Downregulation of SWS cell-autonomously affects surface glia integrity

To test if loss of SWS affects the barrier structure, we analyzed the expression pattern of Coracle (CoraC), which is a major component of septate junctions (Yi et al., 2008). In controls, CoraC is strongly expressed by subperineurial glia cells, shown as a smooth line at the brain surface (Fig. 2A, green arrow). Upon *sws* loss, the CoraC pattern at the brain surface was broken and contained lesions and clumps (Fig. 2B, magenta arrow).

**Figure 2.**
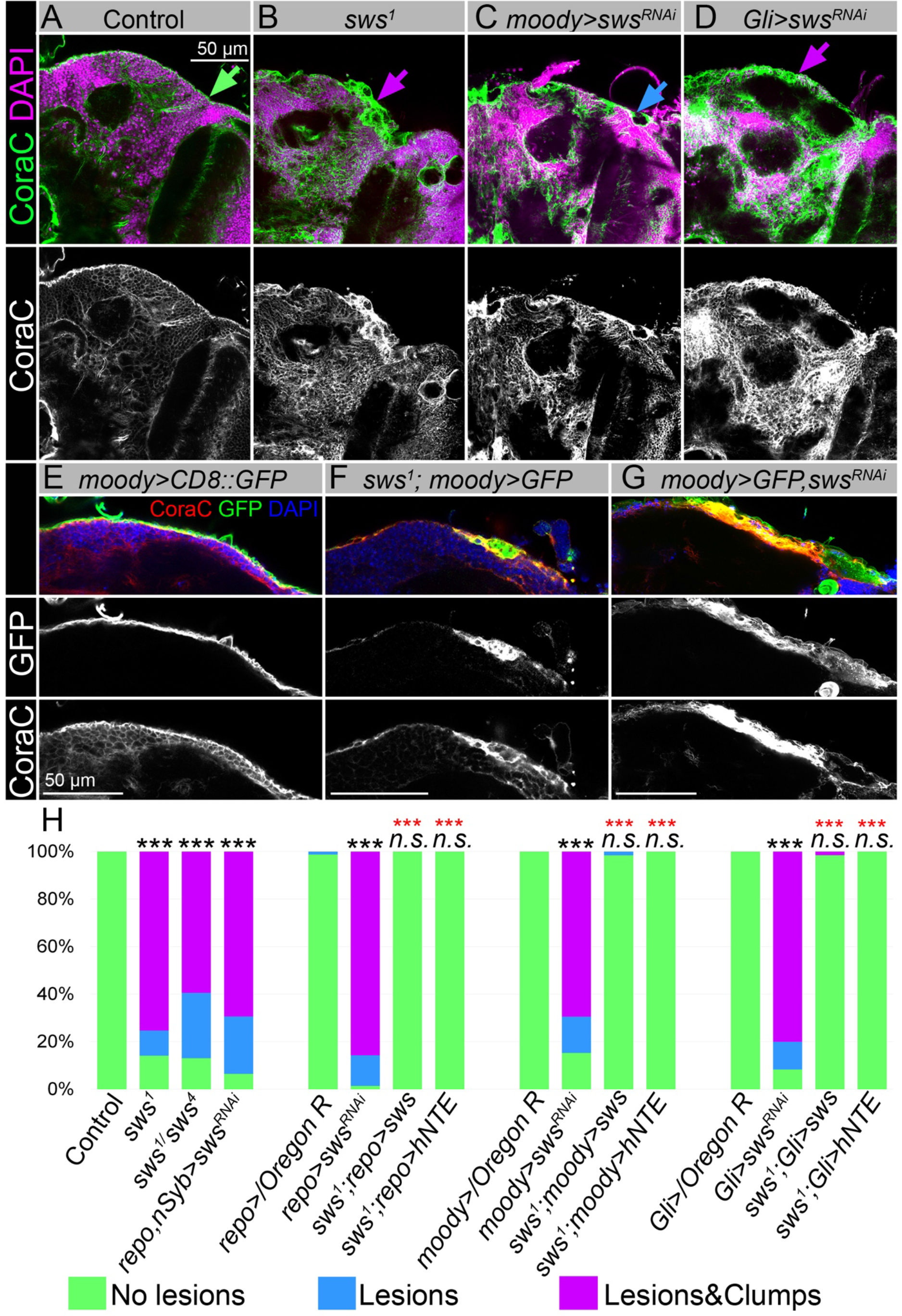
Downregulation of SWS affects surface glia architecture. **A-D.** Adult brains stained with CoraC (green) and DAPI (magenta). **A**. In controls (*Oregon R x white^1118^*), CoraC expression is pictured as the smooth line at the surface of the brain (green arrow). In *sws^1^* mutants (**B**) and in mutants with *sws* downregulation in SPG cells (*moody>sws^RNAi^* and *Gli>sws^RNAi^*, **C** and **D**, respectively), the outer glial cell layer labeled by CoraC is irregular and contains either lesions (blue arrow) or lesions and clumps (magenta arrows). Scale bar: 50 µm. **E-G.** Adult brains stained with CoraC (red), GFP (green) and DAPI (blue) to detect SPG cell membranes marked by co-expression of CoraC and *moody>CD8::GFP* (red+green=yellow). **E.** A smooth line of SPG cell membranes is observed at the surface of control brains (*moody>CD8::GFP*). **F.** In *sws* loss-of-function mutants (*sws^1^; moody>CD8::GFP*), most of the vacuoles formed near the brain surface are filled with the GFP-positive SPG membranes. **G.** Downregulation of *sws* specifically in SPG cells *(moody>sws^RNAi^*) results in the appearance of the same excessive SPG cell membranes inside the brain lesions. Scale bar: 50 µm. **H.** Bar graph shows the percentage of the brain hemispheres with defective brain surface. The percentage of the brain hemispheres with normal brain surface is shown in green, the percentage of the brain hemispheres containing lesions is shown in blue, and the percentage of the brain hemispheres with formed lesions and clumps in the brain surface is shown in purple. Two-way tables and chi-square test were used for statistical analysis. *p < 0.05, **p < 0.005, ***p < 0.001, black asterisks – compared to *Gal4-driver x OR*, red asterisks – compared to *Gal4-driver x UAS-sws^RNAi^*, n of adult brain hemispheres > 43, at least three biological replicates.

Previous characterization of the *sws* loss-of-function mutant showed that SWS deficiency resulted in the formation of membranous glial structures, especially in the lamina cortex (Kretzschmar et al., 1997). Since *sws* is ubiquitously expressed, we specifically downregulated *sws* in the nervous system using the double driver line that allows downregulation of *sws* in glia and neurons (*repo, nSyb-Gal4,* Suppl. Fig. 2C-Cʹ). Since these animals had the same disorganized structure of brain surface as the loss-of-function mutant, we concluded that SWS functions specifically in the nervous system to preserve brain surface structure. Moreover, downregulation of *sws* in all glial cells (*repo>sws^RNAi^*) resulted in the same phenotype. At the same time, upon *sws* downregulation in neurons, we did not observe formation of lesions and clumps in the brain surface, indicating a cell-autonomous function of SWS in glia to maintain BBB organization (Suppl. Fig. 4).

To test if NTE/SWS has a cell-autonomous role in the brain barrier cells, we used already existing subperineurial glia driver lines (*moody-Gal4; UAS-CD8::GFP* and *Gli-Gal4; UAS-CD8::GFP,* Suppl. Fig. 2A-B) and *UAS-sws^RNAi^*(Suppl. Fig. 3). We found that, similar to pan-glial *sws* knockdown, its downregulation specifically in SPG cells caused the formation of lesions and clumps at the brain surface (Fig. 2C-D, blue and magenta arrows). Importantly, expression of *Drosophila* or human NTE in these glia cells rescued this phenotype (Fig. 2H), demonstrating the conserved function of this protein in SPG cells for brain surface formation and possibly maintenance of the brain barrier.

There have been remarkable recent advancements in the field of protein structure prediction, offering valuable tools for exploring three-dimensional structures with unprecedented effectiveness. We used the AlphaFold2 prediction and the PyMol tools (Jumper et al., 2021) to display predicted structure models of the human NTE and *Drosophila* SWS proteins. Both proteins contain a highly conserved patatin-like phospholipase (EST) domain (Suppl. Fig. 5, EST domain in magenta). EST domains in SWS (952-1118) and NTE (981-1147) demonstrated a remarkably high level of confidence, exhibiting helical structures with predicted local distance difference test scores (pLDDT) exceeding 90 (Suppl. Fig. 5). The EST domain exhibits a distinctive architectural pattern comprising three layers of α/β/α structure. Its central region is formed by a six-stranded β-sheet, flanked by α-helices in the front and back. Upon comparing the predicted structures of EST-SWS and EST-NTE, we observed a significant overlap between them (Suppl. Fig. 5). These findings offer additional evidence of the high conservation of functional domains in NTE/SWS and the close relationship between these proteins across different species.

Together, the remarkable similarities observed between human and *Drosophila* SWE/NTE protein structure along with their shared involvement in the formation and maintenance of the brain barrier in *Drosophila* emphasize their close relationship and suggest a conserved function in BBB maintenance.

### Downregulation of SWS results in multilamellar accumulations

Next, we aimed to understand the nature of the SPG phenotype caused by *sws* deficiency. SPG cells have a very specific shape; they are thin and very large. Fewer than fifty SPG cells surround one adult brain hemisphere and a single SPG cell can cover the size of one half of the imaginal disc of the eye, covering an area equivalent to approximately 10,000 epithelial cells (Hartenstein, 2011; Limmer et al., 2014; Silies et al., 2007). Therefore, to better visualize the defects in surface glia organization upon *sws* loss, we introduced *moody-Gal4*; *UAS-CD8::GFP* (*moody>CD8::GFP)* constructs into the *sws^1^*mutant background, which allowed analysis of SPG cell membranes. To our surprise, we observed that almost all lesions that were formed near the brain surface contained membrane material marked by *CD8::GFP* (Fig. 2F). This was in sharp contrast to the control, where SPG membranes formed a distinct GFP-positive line (Fig. 2E). Importantly, the same excessive SPG cell membranes were observed inside the lesions formed upon *sws* downregulation explicitly in SPG cells (Fig. 2G), confirming that SWS is required cell-autonomously in SPG cells for the proper architecture of the surface glia.

Next, we wanted to understand the origin of these excessive membranes observed in *sws*-deficient glial cells. This task appeared to be quite challenging, as SPG cells form a very thin polarized endothelium, not even reaching 1 μm thickness in most areas (Limmer et al., 2014). In addition, SPG cells localize in very close proximity to each other and to neurons, making the analysis of subcellular protein localization challenging. Therefore, to dissect in more detail the *sws*-related phenotype of accumulated SPG membranes inside the lesions on the brain surface, we used an electron microscopy approach.

We found that *sws* mutants showed the formation of various multilamellar bodies in the brain, which were not observed in the control (Fig. 3A-Bʹ). These atypical structures ranged in size from 5 to 15 μm and contained concentrically laminated and multilayered membranes (yellow arrows), lipid droplets (red arrows) and other partially degraded organelles or cytoplasmic constituents. We hypothesized that these inclusions most likely correspond to secondary lysosomes in the phase of digesting endosomal cargo, which are a hallmark of lysosomal storage disorders (LSDs).

**Figure 3.**
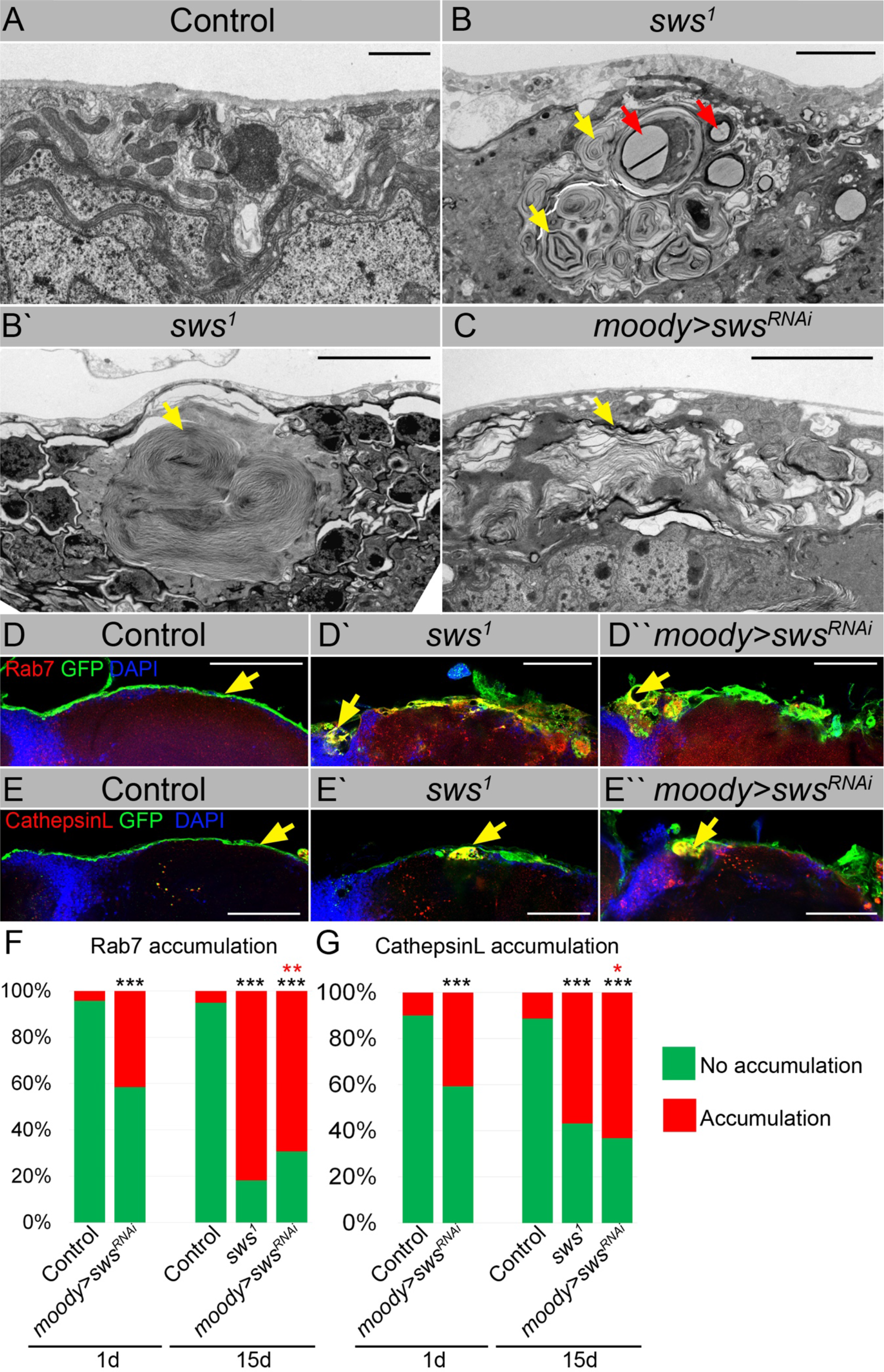
Downregulation of SWS results in intracellular accumulations. **A-C**. Electron microscope images of the surface area of the adult brains. **A.** In controls (*white^1118^*), glia cells that do not contain any abnormal subcellular structures. Scale bar: 1 µm. **B-Bʹ**. *sws^1^*mutant brains have irregular surface and abnormal accumulation of endomembranous structures (yellow arrows) and lipid droplets (red arrows). Scale bar: 5 µm. **C.** *moody>sws^RNAi^* fly brain has same abnormal accumulations of endomembranous structures (yellow arrows) as *sws* mutant. Scale bar: 5 µm. **D-Dʹʹ.** Adult brains stained with Rab7 (red) to detect lysosomes and late endosomes, GFP (*moody>CD8::GFP,* green) to mark SPG cell membranes and DAPI (blue) to mark nuclei. (**D.** A smooth line of SPG cell membranes is observed at the surface of control brains (*moody>CD8::GFP,* green), Rab7 is present in relatively small amounts and evenly dispersed throughout in the brain (red). **Dʹ.** In *sws* loss-of-function mutants (*sws^1^; moody>CD8::GFP*), Rab7-positive structures colocalized with atypical membrane aggregates of GFP-positive SPG membranes (red+green=yellow). **Dʹʹ.** Downregulation of *sws* specifically in SPG cells *(moody> sws^RNAi^*) results in the appearance of the same assemblies in the SPG cells (yellow). Scale bar: 50 µm. **E-Eʹʹ.** Adult brains stained with CathepsinL (red) to detect marks lysosomes, GFP (*moody>CD8::GFP,* green) to mark SPG cell membranes and DAPI (blue) to mark nuclei. **A. E.** A smooth line of SPG cell membranes is observed at the surface of control brains (*moody>CD8::GFP,* green), CathepsinL is present in relatively small amounts in the brain (red). **Dʹ.** In *sws* loss-of-function mutants (*sws^1^; moody>CD8::GFP*), CathepsinL-positive structures colocalized with atypical membrane aggregates of GFP-positive SPG membranes (red+green=yellow). **Eʹʹ.** Downregulation of *sws* specifically in SPG cells *(moody> sws^RNAi^*) results in the appearance of the same assemblies in the SPG cells (yellow). Scale bar: 50 µm. **B. F.** Bar graph shows the percentage of brains with the accumulated Rab7 structures in the brain surface. Two-way tables and chi-square test were used for statistical analysis. *p < 0.05, **p < 0.005, ***p < 0.001, black asterisks – compared to *moody-Gal4 x OR*, red asterisks – compared to 1 day old *moody-Gal4 x UAS-sws^RNAi^*, n of adult brain hemispheres > 44, at least three biological replicates. **C. G.** Bar graph shows the percentage of brains with the accumulated CathepsinL structures at the brain surface. Two-way tables and chi-square test were used for statistical analysis. *p < 0.05, **p < 0.005, ***p < 0.001, black asterisks – compared to *moody-Gal4 x OR*, red asterisks – compared to 1 day old *moody-Gal4 x UAS-sws^RNAi^*, n of adult brain hemispheres > 49, at least three biological replicates.

To authenticate the nature of membranous accumulation in *sws* mutants, we used endosomal and lysosomal markers. Rab7 is a small GTPase that belongs to the Rab family and controls transport to late endocytic compartments such as late endosomes and lysosomes (Guerra and Bucci, 2016). Immunohistochemical analysis demonstrated that in contrast to controls, where Rab7 was present in relatively small and evenly dispersed throughout the brain late endosomes and lysosomes (Fig. 3D, red), in *sws*-deficient brains, accumulation of Rab7-positive compartments was observed. Moreover, Rab7-positive structures colocalized with atypical membrane aggregates of SPG cells (Fig. 3Dʹ, yellow). The same assemblies were observed upon *sws* downregulation in SPG cells (Fig. 3Dʹʹ, yellow).

Rab7 controls biogenesis of lysosomes and clustering and fusion of late endosomes and lysosomes (Feng et al., 2014). Therefore, to support the idea that these abnormal cellular accumulations are of lysosomal origin, we used an additional marker – CathepsinL, which is a key lysosomal proteolytic enzyme expressed in most eukaryotic cells (Xu et al., 2021). We found that *sws* loss or its downregulation in barrier-forming glia cells resulted in the appearance of CathepsinL-positive inclusions that co-localized with GFP-labeled membrane aggregates formed in the mutant SPG cells (Fig. 3E-Eʹʹ, yellow). We conclude that the structures observed upon SWS/NTE deregulation are abnormally enlarged lysosomes.

Next, we quantified the number of brain hemispheres with atypical Rab7- or CathepsinL-positive accumulations. In the control groups, very few (<10%) of the analyzed brains showed accumulation of Rab7 or CathepsinL. However, in mutants with *sws* loss of function and with *sws* SPG-specific downregulation, a significant increase in the frequency of brains containing Rab7- or CathepsinL-positive aggregates was observed (Fig. 3F and 3G). Since *sws*-associated neurodegeneration is age-dependent (Dutta et al., 2016; Kretzschmar et al., 1997; Melentev et al., 2021; Sujkowski et al., 2015; Sunderhaus et al., 2019), we tested if abnormal lysosomes positive for Rab7 and CathepsinL increase with age. Analysis of the brains of 15-day old *sws* downregulation in SPG cells demonstrated ∼2-fold increase in the percentage of brains with lysosomal accumulations at the brain surface in comparison to 1-day old animals (Fig. 3F and 3G). These data demonstrate for the first time that SWS/NTE–associated phenotypes might be additionally characterized by the excessive storage of cellular material in lysosomes that is accelerated by age.

Importantly, similar abnormal buildup of cellular material in lysosomes have been found in hippocampal neuropil (Akassoglou et al., 2004) and spinal axons of NTE-deficient mice (Read et al., 2009). While these structures have not been specifically described as lysosomal defects, the presence of similar dense bodies containing concentrically laminated and multilayered membranes in NTE-deficient mice suggests that, similar to *Drosophila*, SWS/NTE-related phenotypes in mammals may also be associated with excessive storage of cellular material in lysosomes.

### Loss of SWS leads to cell death in brain cells

Lysosomal changes and dysfunction have been involved in the initiation and development of numerous diseases, such as cancer, autoimmune, cardiovascular, neurodegenerative and LSDs (Cao et al., 2021; Hebbar et al., 2017). In particular, LSDs are a group of rare metabolic disorders caused by inherited defects in genes that encode proteins vital for lysosomal homeostasis, such as lysosomal hydrolases or membrane proteins. LSDs often manifest as neurodegenerative disorders. Neurons are rich in lysosomes and are commonly affected by the accumulation of undegraded material. In LSD patients and animal models, cell death has been observed in cell types in the brain other than neurons (Kiselyov et al., 2007). At the same time, not all cells that have increased storage are affected by cell death.

We analyzed if cell death is induced in *sws*-mutant brains and found a dramatic increase of cells positive for the apoptotic marker Cas3 inside mutant brains when compared to controls (Suppl. Fig 6). However, *sws*-mutant SPG cells *per se* did not exhibit an obvious increase in cell death (Suppl. Fig 6), suggesting that surface glia have a cell non-autonomous effect on brain health. Therefore, next, we wanted to understand which other cellular functions are affected by the accumulation of extra material in the lysosomes of SPG cells, resulting in progressive brain degeneration.

### Downregulation of SWS affects brain permeability barrier

The main function of SPG cells is to protect the central nervous system from being exposed to molecules that are harmless to peripheral organs but toxic to brain neurons. SPG cells form a thick polarized endothelium, selective permeability of which is achieved by forming very tight septate junctions that provide structural strength and a barrier that controls the flow of various solutes from outside the brain (Hartenstein, 2011; Limmer et al., 2014; Silies et al., 2007). Since our data show that the expression pattern of a key septate junction protein, Cora, is dramatically perturbed in *sws*-mutant brains (Fig. 2A-B), we decided to test if deregulation of SWS can affect the ability of SPG cells to form a selective permeability barrier.

As a result of abnormal BBB function, the CNS becomes permeable to small molecules such as dextran-coupled dyes. To test BBB permeability, the 10 kDa dextran dye was injected into the abdomen of flies (Fig. 4A). After injection, animals were allowed to recover for at least 12 hours, followed by the dissection and analysis of adult brains. In controls, dextran dye stayed at the outer surface of the brain (Fig. 4B-Bʹ), showing that the BBB is not permeable to 10 kDa dextran. In contrast, the dye was detected inside the brain of *sws^1^* mutants (Fig. 4C, Cʹ). Moreover, the downregulation of *sws* in different types of glial cells also caused increased permeability of brain barrier in more than 80% of the analyzed brains (Fig. 4D). Expression of NTE/SWS in glia in *sws^1^* mutant rescued the organization of the surface glia (Fig. 2H) and partially rescued the barrier phenotype, suggesting that NTE and SWS are important for the BBB integrity in *Drosophila* (Fig. 4D). Taken together, our results demonstrate that SPG cells with SWS deficiency are characterized by defective brain barrier function and lysosomal accumulation of excess cellular material, which includes membranes.

**Figure 4.**
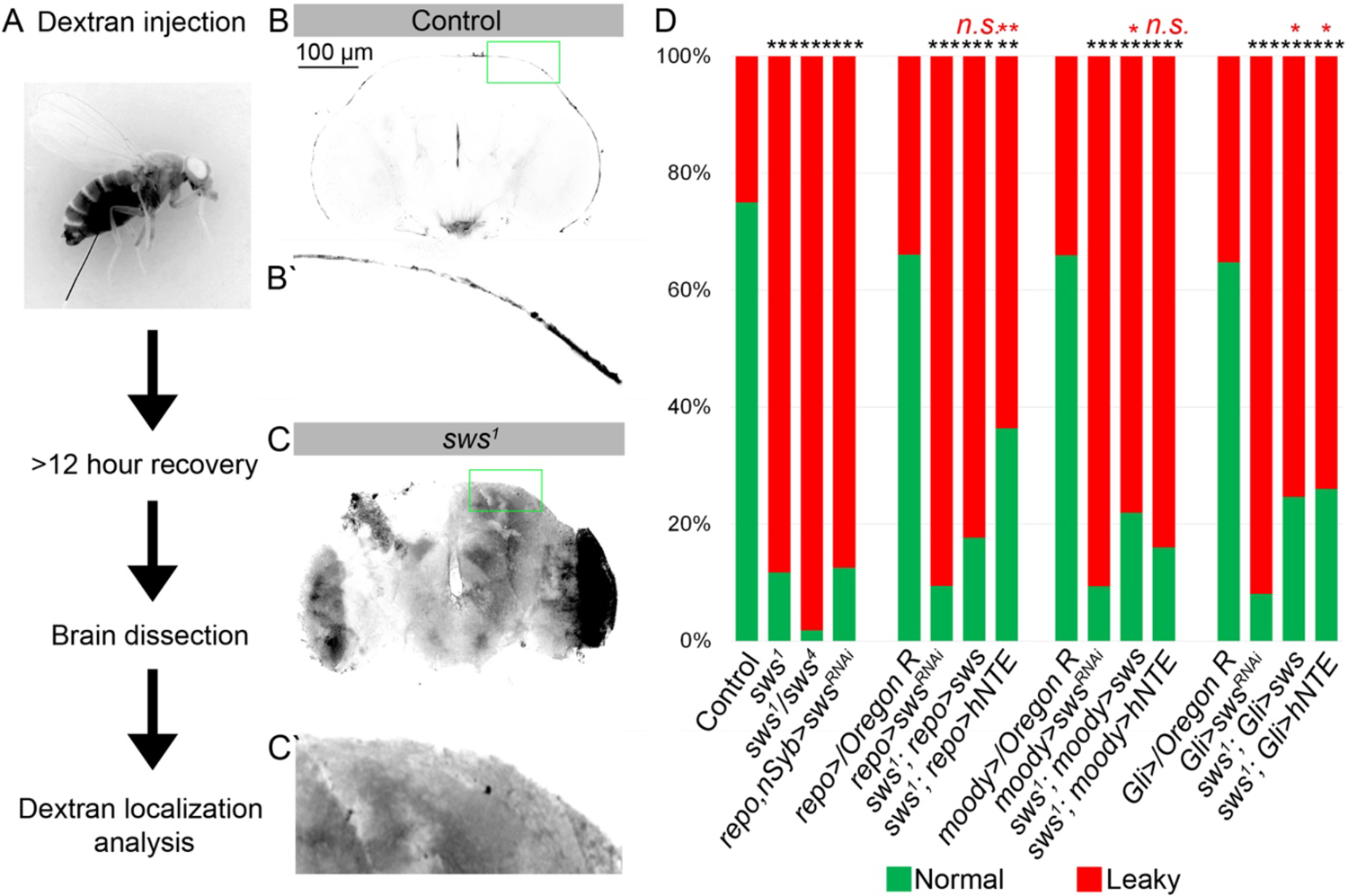
Downregulation of SWS affects brain permeability barrier. **A.** Scheme of 10 kDa dextran dye permeability assay (see also Materials and Methods for a detailed description of the procedure). **B-C.** Localization of dextran dye more than 12 hours after injection in control (*Oregon R*) flies **(B-Bʹ)** and in *sws^1^* mutant **(C-Cʹ)**. Note that dextran dye can be detected in the cells present inside the mutant brain in contrast to control, where dye stays at the outer surface of the brain. Scale bar: 100 µm. **A. D.** Bar graph shows the percentage of the brains with the defective permeability barrier. Two-way tables and chi-square test were used for statistical analysis. *p < 0.05, **p < 0.005, ***p < 0.001, black asterisks – compared to *Gal4-driver x OR*, red asterisks – compared to *Gal4-driver x UAS-sws^RNAi^*, n > 44 of adult brain hemispheres, at least three biological replicates.

Next, we wanted to understand what signaling pathway is activated as a result of leaky brain barrier in *sws* mutants. We treated mutant flies for 14 days with different anti-inflammatory substances and stress suppressors, including apoptosis, ferroptosis, oxidative stress, and ER stress inhibitors and analyzed whether observed glial phenotypes could be suppressed by any medication. We analyzed CoraC expression and compared the frequencies of abnormal brain surface appearance in the drug-treated versus untreated animals (Suppl. Fig. 7A). We revealed that sodium salicylate, a non-steroidal anti-inflammatory drug (NSAID) and rapamycin, which activates autophagy by inhibiting Tor (Xu et al., 2017), showed the best ability to suppress surface glia phenotypes in *sws* mutants (Suppl. Fig. 7B). This indicates that an activated inflammatory response is associated with *sws* deficit.

### moody^ΔC17^ flies with a permeable BBB show glial phenotype similar to sws mutants

A leaky blood-brain barrier allows different toxic substances and bacteria to enter the central nervous system and affect neurons and glial cells, which can lead to cell death and increased inflammation in mammals (Kim et al., 2012). To test if a permeable brain barrier in general is causing inflammation in *Drosophila*, we decided to test if an additional mutant with defective BBB has an increased inflammatory response in the brain. We focused on a *moody^ΔC17^* mutant that has been previously shown to have a defective brain barrier (Bainton et al., 2005).

Firstly, we tested whether the *moody* mutant shows a phenotype similar to that observed in *sws* mutants by analysis of the CoraC expression pattern. We observed that the surface brain layer in *moody* mutants or upon *moody* downregulation in SPG by *moody-Gal4* (*moody>moody^RNAi^*) contained lesions and had an abnormal membrane assembly, resembling CoraC expression pattern in *sws* mutants (compare Suppl. Fig. 8A-C and Fig. 2B, magenta arrows).

Secondly, we analyzed if anti-inflammatory factors can reduce glial phenotypes in *moody* mutants, similar to *sws* mutants. We found that in *moody* mutants, the surface glia phenotype analyzed using CoraC as a marker could also be suppressed by NSAID and rapamycin (Fig. 5A). The fact that anti-inflammatory factors can reduce glial phenotypes in both *sws* and *moody* mutants indicates that inflammation, triggered as a result of a compromised brain barrier, plays a role in a feedback loop that exacerbates the abnormal surface glia organization (Fig. 5D). Next, we tested whether inflammatory pathways are activated in both mutants with permeable barriers.

**Figure 5.**
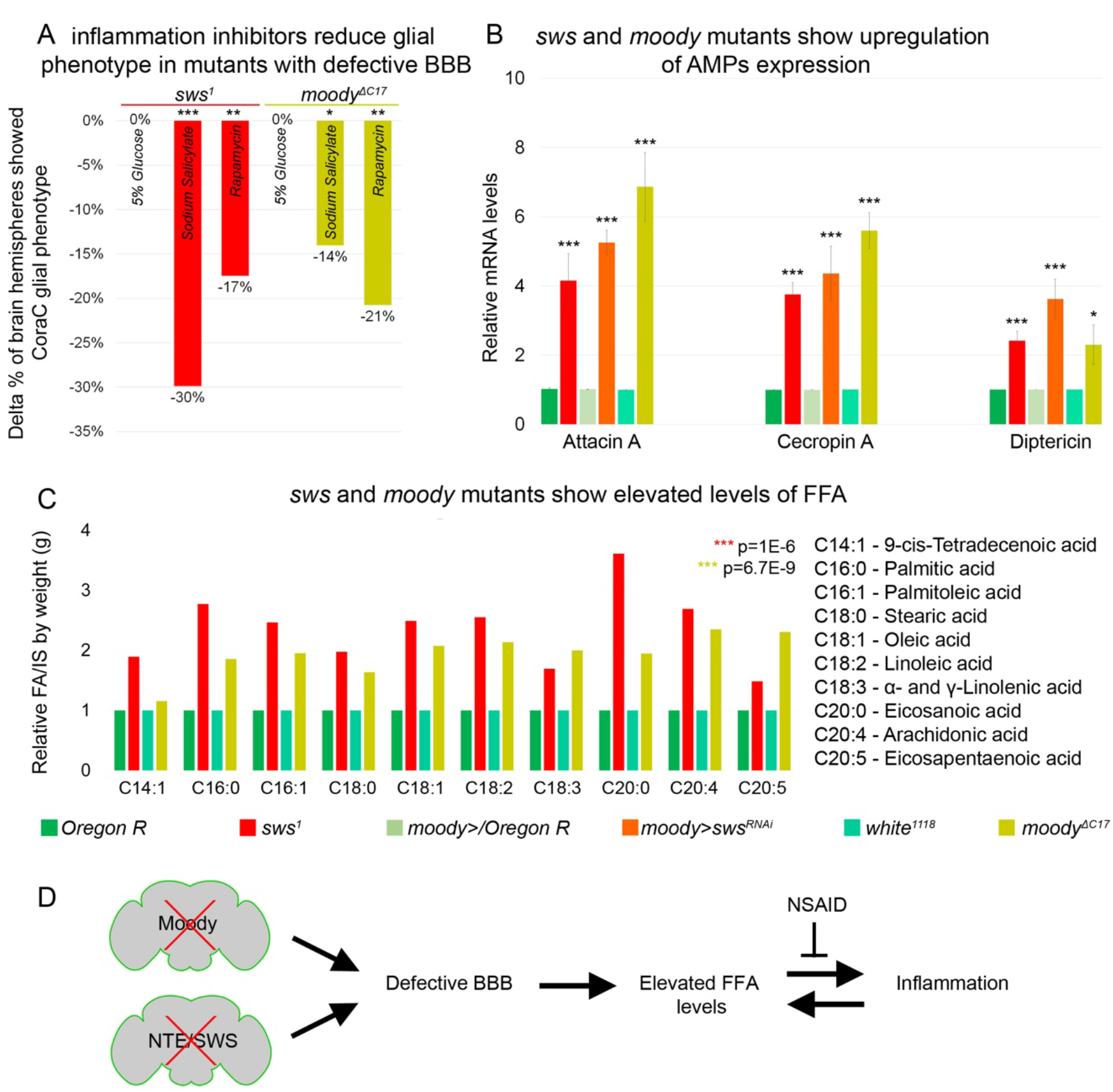
Mutants with defective BBB have increased inflammatory response and elevated levels of FFA. **A.** Bar graph shows the reduction of the percentage of the glial phenotype assayed by CoraC expression pattern in *sws^1^* (red) and *moody* (olive) mutants that were treated with NSAID and rapamycin in comparison to untreated mutants, which suggests that inflammation accelerates surface glia phenotype. Two-way tables and chi-square test were used for statistical analysis (n of adult brain hemispheres > 104, at least three biological replicates). **B.** RT-qPCR analysis of AMPs mRNA levels from relevant controls (green) and *sws^1^* (red), *moody>sws^RNAi^* (orange), and *moody* (olive) mutant fly heads shows significantly upregulated expression of inflammatory response genes: *Attacin A, Cecropin A,* and *Diptericin*. Two-tailed Student’s test was used to test for statistical significance. *Oregon R* vs *sws^1^* mutants: Attacin A: p=5.2E-10, Cecropin A: p=9.1E-15, Diptericin: p=1.3E-6; *moody>/Oregon R* vs *moody>sws^RNAi^*: Attacin A: p=6.9E-11, Cecropin A: p=1.3E-7, Diptericin: p=7.5E-8; *white^1118^*vs *moody* mutants: Attacin A p=1.1E-12, Cecropin A: p=6.8E-13, Diptericin: p=1.3E-6, at least two biological replicates. **C.** GS-MS measurements of free fatty acids (FFA) indicate the relative increase of several free fatty acids in the heads of *sws^1^* (red) and *moody* (olive) mutants compared to relevant controls (*Oregon R* and *white^1118,^* green). One-way Anova test was used for statistical analysis, *p < 0.05, **p < 0.005, ***p < 0.001). **D.** Mutants with defective brain barrier have upregulated innate immunity factors and exhibit elevated levels of free fatty acids involved in mediating the inflammatory response. Treatment with anti-inflammatory agents alleviates BBB phenotypes, suggesting that a signaling loop that links the condition of the brain barrier permeability, lipid metabolism, and inflammation.

### Mutants with defective BBB show upregulation of several innate immunity factors and free fatty acids

The molecular mechanisms of innate immunity between flies and mammals are highly evolutionarily conserved. For example, *Drosophila* Toll and immune deficiency (IMD) pathways are nuclear factor kappa B (NF-κB)-based signaling pathways that share similarities with the Toll-like receptor and tumor necrosis factor receptor 1 signaling pathways in mammals (Cao et al., 2013; Kounatidis and Chtarbanova, 2018; Kounatidis et al., 2017; Pavlidaki et al., 2022). It has been previously shown that in glial cells, activation of the IMD pathway results in phosphorylation of the NF-κB transcription factor Relish, which is translocated to the nucleus to induce expression of the antimicrobial peptides (AMPs) Attacin A, Cecropin A, and Diptericin (Kounatidis and Chtarbanova, 2018; Winkler et al., 2021). We performed qPCR analysis and measured the mRNA levels of these AMPs in heads of *sws* and *moody* loss-of-function mutants and flies that had *sws* downregulation only in SPG cells (*moody>sws^RNAi^*). We found that the mRNA levels of all three AMPs were significantly upregulated in mutants in comparison to relevant controls (Fig. 5B). These data demonstrate that both mutants with defective BBB exhibit an increased inflammatory response and that surface glia-specific downregulation of *sws* is sufficient to induce inflammation.

In addition, polyunsaturated fatty acids have been shown to play a key role in inflammatory processes. Their oxygenated products, called eicosanoids, induce and regulate inflammation via G-protein coupled receptor (GPCR) signaling pathways (Stanley and Kim, 2018). To find out whether levels of polyunsaturated and saturated fatty acids are changed, we measured levels of free fatty acids from accurately weighed heads of control flies and mutants with defective BBB (*sws^1^* and *moody ^ΔC17^*). Free fatty acids were measured by gas chromatography-mass spectrometry (GC-MS) as described recently (von Hanstein et al., 2023). We found that both mutants with defective BBB show upregulated levels of linoleic acid, α- and γ-linolenic acid, eicosanoic acid, arachidonic acid, and eicosapentaenoic acid (EPA) when compared to controls (Fig. 5C). Additionally, levels of other free fatty acids involved in inflammatory response, 9-cis-tetradecenoic acid, palmitic acid, palmitoleic acid, stearic acid, and oleic acid (Korbecki and Bajdak-Rusinek, 2019; Miao et al., 2015) were elevated upon *sws* or *moody* loss (Fig. 5C). These data show that the inflammatory response accompanied by the accumulation of free fatty acids is induced in both mutants with leaky BBB.

### sws and moody mutants have distinct surface glia phenotypes

However, while both *sws* and *moody* mutants have defective BBB, the nature of these mutations and their involvement in cellular processes are very different. Moody is a GPCR that is expressed in SPGs and localizes to the sites of septate junction formation (Babatz et al., 2018). Its cellular function is to control continued cell growth of SPG by differentially regulating actomyosin contractility and septate junction organization (Li et al., 2021b). SWS is a transmembrane ER protein that hydrolyzes phosphatidylcholine and binds to and inhibits the C3 catalytic subunit of protein kinase A (Bettencourt da Cruz et al., 2008). To understand how such different mutations could result in similar outcomes, we first analyzed if *moody* loss would result in lysosomal material accumulation. Electron microscopy analyses demonstrated that unlike in *sws* mutant brains, no intracellular accumulations with extra cellular material were observed upon *moody* loss (compare Fig 3. B-Bʹ and Suppl. Fig. 8F). Furthermore, no accumulation of endosomal-lysosomal pathway components such as Rab7 were detected within SPG cells of *moody* mutants (Suppl. Fig. 9A, B). At the same time, as previously described (Babatz et al., 2018; Schwabe et al., 2005), we observed that in the absence of *moody*, septate junctions were formed, but they were disorganized (Suppl. Fig. 8F, arrow).

We compared in greater detail the septate junction organization in both mutants using a molecular component of septate junctions, Neurexin IV (Nrx-IV). In comparison to the wild type, upon *sws* loss, septate junctions were not properly assembled and exhibited irregular clumps and disruptions (Fig. 6A, C). In contrast, the *moody* mutant exhibited a frayed septate junction phenotype (Fig. 6B). Since Moody coordinates the continuous organization of junctional strands in an F-actin-dependent manner, as a result of its loss, septate junction strands fail to extend properly during cell growth (Fig. 6C). While the role of Moody in septate junction formation is understood (Babatz et al., 2018), the mechanism by which SWS may be involved in this process is unclear.

**Figure 6.**
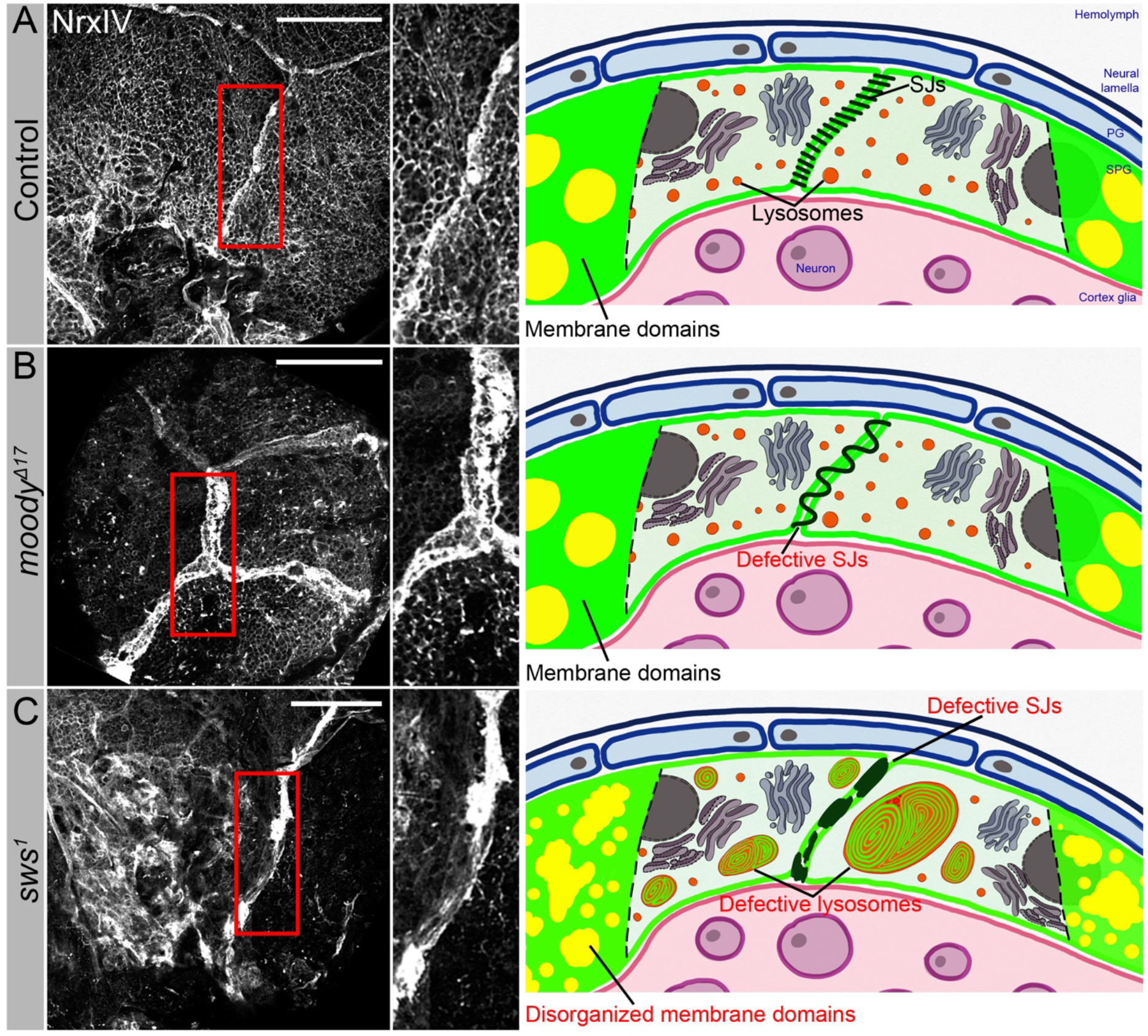
Septate junction and membrane domain organization in mutants with defective brain permeability barrier. **A-C.** Adult brains stained with a septate junction marker NrxIV (white). Scale bar: 50 µm. In control (*Oregon R*) brain, septate junctions formed by SPG glia are depicted as condensed and distinct strand **(A)**. The scheme depicts the intact blood-brain barrier (BBB) formed by perineurial glia (PG) and subperineurial glia (SPG). The SPG cells establish well-formed septate junctions (SJs) and exhibit organized membrane domains. Furthermore, the lysosomes are fully functional. In *moody^ΔC17^* mutants, due to SPG membrane overgrowth, septate junctions are frayed **(B)**. The scheme illustrates a defective BBB where the proper extension of septate junction strands during cell growth is impaired, resulting in increased permeability. However, despite this issue, the membrane domains remain well-formed, and the lysosomes within the barrier continue to function effectively. In *sws^1^* mutants, septate junctions and membrane domains are not properly organized **(C)**. By analyzing SPG membranes in *sws* mutants, abnormal clustering of septate junction (SJ) proteins and disorganized membrane domains are observed. Furthermore, *sws*-deficient brains exhibit excessive storage of cellular material within lysosomes. The scheme shows that SWS-related lipid dysregulation is accompanied by dysfunctional lysosomes, impaired distribution of cell junction proteins, and disrupted organization of membrane domains in surface glia.

Cell junctions are a special type of plasma membrane domain whose transmembrane proteins form a complex, mechanically stable multiprotein structure (Giepmans and van Ijzendoorn, 2009). The lipid component of cell junctions exhibits a typical membrane raft structure (Lee et al., 2008; Muhlig-Versen et al., 2005; Nusrat et al., 2000; Shigetomi et al., 2023; Simons and Vaz, 2004). The main feature of membrane rafts is that they contain an enriched fraction of cholesterol and sphingolipids and are able to dynamically orchestrate specific membrane proteins involved in cell adhesion, signal transduction, protein transport, pathogen entry into the cell, *etc.* Since SWS regulates lipid membrane homeostasis, we hypothesized that SWS influences the composition of membrane rafts. Analysis of SPG membranes in *sws*-deficient brains shows abnormal clustering of septate junction proteins and disorganized membrane domains, implying that SWS phospholipase plays a role in organizing SPG membrane architecture (Fig. 6C).

In summary, our data show that the phospholipase NTE/SWS plays a crucial role in lysosome biogenesis and organization of the architectural framework of BBB membranes. We propose that since NTE/SWS regulates lipid membrane homeostasis, is loss results in the disruption of membrane rafts, which includes septate junctions, leading to brain barrier permeability. As a result, the inflammatory response associated with the accumulation of free fatty acids is activated in mutant brains, leading to progressive neurodegeneration.

## DISCUSSION

The physiological functions of the BBB, maintaining and protecting the homeostasis of the CNS, are evolutionarily conserved across species (Bundgaard and Abbott, 2008). Even though there is already plenty of evidence connecting BBB dysfunction to neurodegenerative diseases, the underlying mechanism is not fully understood. The BBB is formed by microvascular endothelial cells lining the cerebral capillaries penetrating the brain and spinal cord of most mammals and other organisms with a well-developed CNS (Kadry et al., 2020). Interestingly, NTE is highly expressed not only in the nervous system but also in endothelial cells, suggesting that BBB might be affected upon NTE-associated neurodegenerations (The Human Protein Atlas - https://www.proteinatlas.org/ENSG00000032444-PNPLA6/single+cell+type).

Here we made an intriguing discovery regarding the presence of NTE/SWS in the surface glia responsible for forming the blood-brain barrier (BBB), where it plays a crucial role in ensuring the selective permeability of the BBB and the proper organization of surface glia. Moreover, here we discovered that NTE/SWS-associated neurodegeneration is accompanied by abnormal membrane accumulation within defective lysosomes, indicating importance of NTE/SWS in proper function of lysosomes. It has been demonstrated for some LSDs, for example, Krabbe’s disease, to be pathologically characterized by rapidly progressive demyelination of the central nervous system and peripheral nervous system and accumulation of macrophages in the demyelinating lesions (Kondo et al., 2005). Considering that NTE/SWS is involved in the maturation of non-myelinating Schwann cells during development and de-/remyelination after neuronal injury (McFerrin et al., 2017), suggesting that lysosomal function of NTE/SWS might be essential for proper myelination in vertebrates. Interestingly, we found that loss of *sws* or its downregulation in barrier-forming glia led to accumulations of Rab7 and CathepsinL in these cells, demonstrating that SWS/NTE – associated neuropathies might be additionally characterized by excessive storage of cellular material in lysosomes. Importantly, neuroinflammation has been reported in several LSDs. The most abundant lysosomal proteases, Cathepsins have been shown to contribute to neuroinflammation as well as to induce neuronal apoptosis (Tschopp and Schroder, 2010).

Over the past few years, there has been a growing appreciation of the organizing principle in cell membranes, especially within the plasma membrane, where such domains are often referred to as “lipid rafts”. Such lipid rafts were defined as transient, relatively ordered membrane domains, the formation of which is driven by lipid–lipid and lipid– protein interactions (Sezgin et al., 2017). Previously, it has been demonstrated that NTE/SWS is crucial for membrane lipid homeostasis, and *sws* mutants exhibit increased levels of phosphatidylcholine (Muhlig-Versen et al., 2005). Phosphatidylcholine, a key component of most organellar membranes, possesses an amphiphilic nature, enabling it to energetically self-assemble into continuous bilayers(Yang et al., 2018). This ability to spontaneously self-organize can explain the appearance of multilayered membrane structures in the lysosomes of sws mutants. Furthermore, phosphatidylcholine plays a vital role in generating spontaneous curvature, essential for membrane bending and tubulation in vesicular transport processes within the cell (Epand and Epand, 1994). Therefore, abnormal levels of phosphatidylcholine may impact the lysosome fission and fusion steps, leading to the accumulation of defective lysosomes in *sws* mutants. Since lysosomes are involved in lipid catabolism and transport, disruptions in their function can additionally affect cellular lipid homeostasis (Thelen and Zoncu, 2017). Consequently, alterations in lipid composition due to abnormal NTE/SWS phospholipase function and defective lysosomes in *sws*-mutant cells could affect the constitution of the plasma membrane and its ability to form lipid-driven membrane rafts. Lipid rafts are characterized by the clustering of specific membrane lipids through spontaneous separation of glycolipids, sphingolipids, and cholesterol in a liquid-ordered phase (Grassi et al., 2020). Their assembly dynamics depend on the relative availability of different lipids and membrane proteins (Simons and Vaz, 2004). Lipid rafts play significant roles in multiple cellular processes, including signaling transduction (Sezgin et al., 2017). Interestingly, tight junctions are considered as raft-like membrane compartments (Nusrat et al., 2000), as they represent membrane microdomains crucial for the spatial organization of cell junctions and regulation of paracellular permeability (Lee et al., 2008; Shigetomi et al., 2023). Therefore, we propose that abnormal organization of tight junctions in the SPG cells of *sws* mutants is caused by abnormal organization of plasma membrane domains.

Lysosomes play an essential role in the breakdown and recycling of intracellular and extracellular material, including lipids, proteins, nucleic acids, and carbohydrates. Any dysfunction of lysosomal system components has catastrophic effects and leads to a variety of fatal diseases (Udayar et al., 2022). LSDs are often linked to changes in plasma membrane lipid content and lipid raft stoichiometry (Domon et al., 2011; Vainio et al., 2005), inflammation (DiRosario et al., 2009; Seehafer et al., 2011) and ER stress responses (Kim et al., 2006; Tessitore et al., 2004). In the past few years, treatments for LSDs were only able to deal with signs and symptoms of the disorders. One possible approach is to identify an available source for the deficient enzyme using therapeutic methods such as bone marrow transplantation (BMT), enzyme replacement therapy (ERT), substrate reduction therapy (SRT), chemical chaperone therapy (CCT), and gene therapy. At the present time, such strategies are aimed at relieving the severity of symptoms or delaying the disease’s progression, yet do not provide a complete cure (Sheth and Nair, 2020). However, since we and others (Sujkowski et al., 2015) have shown that overexpression of human NTE can ameliorate mutant phenotype, it can be speculated that, depending on the causative mutation, ERT might be an option as treatment of NTE/SWS-related disorders.

It has been demonstrated that the ER establishes contacts between its tubules and late endosomes (LEs)/lysosomes, visualized in unpolarized cells as well as in neurons derived from brain tissue. Moreover, disruption of ER tubules causes accumulation of enlarged and less-motile mature lysosomes in the soma, suggesting that ER shape and proper function orchestrate axonal late endosome/lysosome availability in neurons (Ozkan et al., 2021; Wu et al., 2017). Considering the ER localization of NTE/SWS in the cell, we propose that abnormal lipid composition in the membrane upon *sws* loss has a significant effect on lysosome structure and functions. Furthermore, ER forms contact sites with plasma membrane through vesicle-associated membrane protein (VAMP)-associated protein VAP (Li et al., 2021a). Loss of VAP results in neurodegeneration, such as sporadic amyotrophic lateral sclerosis or Parkinson’s disease (Anagnostou et al., 2010; Kun-Rodrigues et al., 2015). Mitochondria-ER contact sites play a crucial role in many vital cellular homoeostatic functions, including mitochondrial quality control, lipid metabolism, calcium homeostasis, unfolded protein response, and ER stress. Disruptions in these functions are commonly observed in neurodegenerative disorders like Parkinson’s disease, Alzheimer’s disease, and amyotrophic lateral sclerosis (Wilson and Metzakopian, 2021). Interestingly, knockdown of *sws* in neurons reduces mitochondria number in the brain and in wing axons (Melentev et al., 2021). SWS-deficient animals show activation of ER stress response, characterized by elevated levels of GRP78 chaperone and increased splicing of XBP, an ER transcription factor that triggers transcriptional ER stress responses. Neuronal overexpressing XBP1 and treating flies with tauroursodeoxycholic acid (TUDCA), a chemical known to attenuate ER stress-mediated cell death, alleviated locomotor deficits and neurodegeneration in *sws* mutants assayed by vacuolization area (Sunderhaus et al., 2019). Reduced levels of Sarco/Endoplasmic Reticulum Ca^2+^ ATPase (SERCA) observed in *sws* mutants were linked to disrupted lipid compositions as well. Promoting cytoprotective ER stress pathways may provide therapeutic relief for NTE-related neurodegeneration and motor symptoms (Sunderhaus et al., 2019).

Moreover, we found that BBB disruption is accompanied by elevated levels of free fatty acids, involved in multiple extremely important biological processes. Fatty acids are locally produced in the endothelium and later are transported inside the brain across the BBB (Pifferi et al., 2021). We discovered that *Drosophila* mutants with leaky BBB showed upregulated levels of such fatty acids as palmitoleic, oleic, linoleic, linolenic, arachidonic, and eicosapentaenoic acids, suggesting abnormal metabolism of unsaturated fatty acids upon barrier dysfunction. In particular, *sws* loss results in increased levels of some saturated free fatty acids (FFAs), including palmitic and stearic acids. FFAs or non-esterified fatty acids, are known to be significant sources of ROS, which lead to the event of oxidative stress (Soardo et al., 2011), resulting in lipotoxicity associated with ER stress, calcium dysregulation, mitochondrial dysfunction, and cell death (Ly et al., 2017). Previously it has been demonstrated ROS accumulation and activated ER stress response upon *sws* loss in neurons and glia (Melentev et al., 2021; Ryabova et al., 2021; Sunderhaus et al., 2019), which might be a result of increased levels of FFAs. In addition, neuronal *sws* knockdown results in the upregulation of antioxidant defense genes (Melentev et al., 2021). We found that BBB breakdown is accompanied by abnormal fatty acids metabolism, and rapamycin can suppress the abnormal glial phenotype formed in BBB *Drosophila* mutants. Interestingly, saturated FFAs have been shown to lead to target of rapamycin (mTOR) complex 1 activation and cell apoptosis in podocytes (Yasuda et al., 2014). Moreover, rapamycin significantly diminishes FFA-induced podocyte apoptosis (Yasuda et al., 2014), supporting its potential ability to suppress possible outcomes of FFA upregulation in the *Drosophila* brain, thus improving glial phenotype in mutants with BBB breakdown.

Polyunsaturated fatty acids (PUFAs) are known to be primary precursors of lipid mediators that are abundant immunomodulators (Kwon et al., 2020). Lipid mediators are signaling molecules, such as eicosanoids, and are implicated in inflammation. More recently, lipid molecules that are pro-inflammatory, and those involved in the resolution of inflammation have become important targets of therapeutic intervention in chronic inflammatory conditions. According to published research, PUFAs metabolism was additionally associated with Alzheimer’s disease and dementia (Rao et al., 2017; van der Lee et al., 2018). The focus of particular interest has recently been on the PUFAs’ involvement in the continued inflammatory response because, in contrast to acute inflammation, chronic inflammatory processes within the central nervous system are crucial for the development of brain pathologies (Funk, 2001; Regulska et al., 2021). In our study, we found that brain permeability barrier breakdown is accompanied by abnormal fatty acids metabolism and that an aspirin analogue - a non-steroidal anti-inflammatory drug (NSAID) - showed the best ability to suppress abnormal glial phenotype, indicating that activated inflammatory response possibly plays an important role in maintaining a healthy brain barrier. Thus, feedback signaling loop exists between the condition of the brain permeability barrier, lipid metabolism, and the extent of inflammation.

According to the World Health Organization (WHO), the current decade is considered the Decade of Healthy Aging. As the speed of population aging is accelerating worldwide, the proportion of older people will increase from one in eight people aged 60 years or over in 2017 to one in six by 2030 and one in five by 2050 (Keating, 2022; Rudnicka et al., 2020). Globally, there is a little evidence that older people today are in better health than previous generations (https://www.who.int/home/cms-decommissioning). If people who enter extended age of life are in good health, they will continue to participate and be an integral part of families and communities and will strengthen societies; however, if the added years are dominated by poor health, social isolation or dependency on care, the implications for older people and for society are much more negative. Therefore, aging of the world population has become one of the most important demographic problems/challenges of modern society. Moreover, the global strategy on aging and health of the older population includes not only treating but also preventing some of the world’s leading age-related diseases using biomarkers as indicators of any aspects of health change (Crimmins et al., 2008). Unfortunately, most neurodegenerative diseases in humans currently have no cure, and only palliative care is available. Current research is primarily focused on promoting the development of therapies that can prevent the onset of a number of age-related neurodegenerative diseases. Specific and effective treatments are urgently needed. However, their advance hinges upon a deeper understanding of the molecular mechanisms underlying progressive neurodegeneration. Understanding the molecular mechanisms of inflammaging activated by abnormal fatty acid metabolism and testing new and available drugs in a model organism such as *Drosophila* may help us to promote the use of anti-inflammatory therapy and dietary supplements for neurodegeneration and get closer to preventing and curing the diseases that lead to malfunctions in the aged brain.

## MATERIALS AND METHODS

### Drosophila stocks

Fly stocks were maintained at 25°C on a standard cornmeal-agar diet in a controlled environment (constant humidity and light-dark cycle). As controls *OregonR* and *w^1118^* lines were used. The *sws^1^*mutant and the *UAS-sws* lines were gifts from Doris Kretzschmar (Kretzschmar et al., 1997). To obtain *sws* transheterozygotes, *sws^1^*and *sws^4^,* obtained from Bloomington Drosophila Stock Center (BDSC 28121), mutant alleles were used. To express transgenes in a *sws*-dependent manner, a *sws* driver line (*sws-GAL4*), obtained from the Kyoto Stock Center (104592), was used. To define an expression pattern of the driver lines, a *UAS-nlsLacZ, UAS-CD8::GFP* transgenic line, kindly donated by Frank Hirth, was used. To induce human NTE gene expression, a *UAS-hNTE* transgenic line (kindly donated by Robert Wessells) was used. To downregulate *sws* expression*, UAS-sws^RNAi^* (BDSC 61338) was used. Glia-specific Gal4 driver lines - *repo-Gal4, UAS-CD8::GFP/TM6B*, *Gliotactin-GAL4, UAS-CD8::GFP* and *moody-GAL4, UAS-CD8::GFP* - were gifts from Mikael Simons. A neuronal Gal4 driver, *nSyb-GAL4* was obtained from BDSC (BDSC 51945). In addition, to phenocopy *sws* loss-of-function in the nervous system, a double driver line was generated (*repo-Gal4, nSyb-Gal4, UAS-CD8::GFP/TM6B, Sb),* which allowed the expression of the transgenes in both neuronal and glial cells. The *moody ^ΔC17^* mutant was a gift from Christian Klämbt. To downregulate *moody* expression, *UAS-moody^RNAi^* (BDSC 66326) was used.

### Immunohistochemistry

Fly brains of 15-day-old animals were dissected in 1x Phosphate Buffered Saline (1x PBS) and then fixed in 4% formaldehyde diluted in 1x PBS for 20 minutes at room temperature. Next, brains were washed with PBT (0.2% Triton X-100 in 1x PBS) 4 times, followed by block with PBTB (2 g/l Bovine Serum Albumin, 5% Normal Goat Serum, 0.5 g/l sodium azide) for one hour at room temperature and then incubated at 4°C in with primary antibodies diluted in PBTB on nutator overnight. The following day, samples were washed with 1x PBT four times followed by block for 1h and 2h incubation with secondary antibodies at room temperature. Next, samples were washed 4 times with PBT (one of the washes contained DAPI to mark nuclei). Lastly, medium (70% glycerol, 3% n-propyl gallate in 1x PBS) was added to samples for later mounting on the slides. The following primary antibodies were used: mouse anti-Repo (1:50), mouse anti-CoraC (1:50), mouse anti-Elav (1:50), and mouse anti-Rab7 (1:50), rat anti-DE-Cadherin (1:50) from the Developmental Studies Hybridoma Bank (DSHB); chicken anti-GFP (#ab13970, 1:1000) from Abcam; mouse Anti-β-Galactosidase (#Z3781, 1:200) from Promega; rabbit anti-SWS (1:1000 from Doris Kretzschmar); mouse anti-CathepsinL (#1515-CY-010, 1:400) from R&D Systems; rabbit anti-Cas3 (#9662, 1:1000) from Cell Signaling; and rabbit anti-NrxIV (1:1000 from Christian Klämbt). The following secondary antibodies were used: goat anti-chicken Alexa 488 (1:500), goat anti-rat Alexa 488 (1:500), goat anti-rat Alexa 647 (1:500), goat anti-rabbit Alexa 488 (1:500), and goat anti-rabbit Alex 568 (1:500) from Thermo Fisher Scientific; goat anti-mouse IgG2a Cy3 (1:400), goat anti-mouse IgG1 647, and goat anti-mouse IgG1 Cy3 (1:500) from Jackson ImmunoResearch Laboratory. For visualization of cell nuclei, DAPI dye was used (1:1000, Sigma). Samples were analyzed using a confocal microscope (Zeiss LSM 700). For making figures, Adobe Photoshop software was used.

### RNA preparation and real-time quantitative PCR (RT-qPCR)

Total RNA was extracted from 15-day-old fly brains using Trizol reagent (Invitrogen) following the manufacturer’s protocol. To detect *sws* mRNA, the forward and reverse primers AGACATACGCCGTGAATACCG and GCGACGACTGTGTGGACTTG, respectively were used. To detect expression of innate immunity factors, the forward and reverse primers showed in Supplementary Table 5 were used. As an endogenous control for qPCR reactions, Ribosomal Protein L32 (RpL32) with the following forward and reverse primers AAGATGACCATCCGCCCAGC and GTCGATACCCTTGGGCTTGC, respectively, was used. All reactions were run at least in triplicate with appropriate blank controls. The threshold cycle (CT) was defined as the fractional cycle number at which the fluorescence passes a fixed threshold. The ΔCT value was determined by subtracting the average RpL32 mRNA CT value from the average tested CT value of target mRNA, correspondingly. The ΔΔCT value was calculated by subtracting the ΔCT of the control sample from the ΔCT of the experimental sample. The relative amounts of miRNAs or target mRNA is then determined using the expression 2^−ΔΔCT^.

### Permeability assay

Flies were injected into the abdomen with a solution containing 10 kDa dextran dye labeled with Texas Red (#D1864) from Molecular Probes. Flies were then allowed to recover for more than 12 hours before the dissection, followed by the analysis for dextran dye presence in the brain. Fly heads of 15-day-old animals were dissected in 1x Phosphate Buffered Saline (1x PBS) and then fixed in 4% formaldehyde diluted in 1x PBS for 1 hour at room temperature. Then fly brains were dissected in 1x Phosphate Buffered Saline (1x PBS) and fixed in 4% formaldehyde diluted in 1x PBS for 20 minutes at room temperature. Next, brains were washed with PBT (0.2% Triton X-100 in 1x PBS) 4 times, followed by block with PBTB (2 g/l Bovine Serum Albumin, 5% Normal Goat Serum, 0.5 g/l sodium azide) for one hour at room temperature and then washed 2 times with PBT (one of the washes contained DAPI to mark nuclei). Lastly, medium (70% glycerol, 3% n-propyl gallate in 1x PBS) was added to samples for later mounting on the slides.

### In vivo Drosophila treatments

TUDCA (#580549), 4-PBA (#567616), Valsartan (#PHR1315), Fenofibrate (#F6020), Sodium Salicylate (#S3007), Rapamycin (#R0395), Deferoxamine mesylate salt (#D9533), Liproxstatin-1 (#SML1414), and Sphingosine (#860025P) from Sigma Aldrich were added to 5% Glucose solution at a final concentration showed by Supplementary Table 7. Then, these glucose-dissolved components were fed by micropipettes to the flies that were kept for 14 days on a diet food without any sugar. To visualize the uptake of chemicals, solutions were also colored by 2, 5% w/v of Brilliant Blue (80717).

### Extraction and derivatization of free fatty acids from flies

Accurately weighed heads of 15-day-old flies were treated with 1000-µL aliquots of acetonitrile in autosampler glass vials (1.8 mL), the samples were sealed and vortexed several times and then stored in a refrigerator overnight (4°C). Next day, the samples were wormed up to room temperature and centrifuged (10 min, 3345xg, 4°C). Aliquots (950 µL) of the clear supernatants were decanted carefully transferred to autosampler glass vials (1.8 mL). The samples were spiked with 10-µL aliquots of a 1000-µM stock solution of sterculic acid (C19H34O2; 10 nmol; 8-cyclopropen-octadecenoic acid corresponds to C19:1”) which served as the internal standard (IS) for all free fatty acids. The solvent was evaporated entirely under a stream of nitrogen gas. The solid residues were reconstituted in anhydrous acetonitrile (100 µL). Then, 10 µL Hünig base (*N,N*-diisopropylethylamine) and 10 µL 33 vol% pentafluorobenzyl (PFB) bromide in anhydrous acetonitrile were added. Subsequently, the free fatty acids were derivatized by heating for 60 min at 30°C to generate the PFB esters of the free fatty acids. Solvents and reagents were evaporated to dryness under a stream of nitrogen gas. The residues were treated with 1000-µL aliquots of toluene and the derivatives were extracted by vortex-mixing for 120 s. After centrifugation (10 min, 3345xg, 4°C), 300-µL aliquots of the clear and colorless supernatants were transferred into microvials placed in autosampler glass vials (1.8 mL) for GC-MS analysis. A standard control sample containing 1 mL acetonitrile, 1 µL 10 mM arachidonic acid (C20:4, 10 nmol) and 10 µL 1 mM IS (10 nmol) was derivatized as described above for the fly samples after were evaporation to dryness under a stream of nitrogen gas. After centrifugation (10 min, 3345xg, 4°C), 100-µL of the clear and colorless supernatant were transferred into an autosampler glass vial (1.8 mL), diluted with toluene (1:10, v/v) and subjected to GC-MS analysis as described below.

### GC-MS analysis of free fatty acids from flies

GC-MS analyses were performed on a GC-MS apparatus consisting of a single quadrupole mass spectrometer model ISQ, a Trace 1210 series gas chromatograph and an AS1310 autosampler from ThermoFisher (Dreieich, Germany). A fused-silica capillary column Optima 17 (15 m length, 0.25 mm I.D., 0.25 µm film thickness) from Macherey-Nagel (Düren, Germany) was used. Aliquots of 1 µL were injected in the splitless mode. Injector temperature was kept at 280°C. Helium was used as the carrier gas at a constant flow rate of 1.0 mL/min. The oven temperature was held at 40°C for 0.5 min and ramped to 210°C at a rate of 15°C/min, and then to 320°C at a rate of 35°C/min. Interface and ion-source temperatures were set to 300°C and 250°C, respectively. Electron energy was 70 eV and electron current 50 µA. Methane (constant flow rate of 2.4 mL/min) was used as the reactant gas for negative-ion chemical ionization (NICI). The electron multiplier voltage was set to 1300 V. Authentic commercially available reference compounds were used to determine the retention times of the derivatives and to generate their mass spectra. The selected ions [M−PFB] − with mass-to-charge (*m/z*) ratios and retention times of the derivatives are summarized in Supplementary Table 6b. Quantitative measurements were performed by selected-ion monitoring (SIM) of the ions listed in Supplementary Table 6b with a dwell time of 50 ms and SIM width of 0.5 amu for each ion in three window ranges. The results of the GC-MS analyses of the control standard sample that contained 10 nmol arachidonic acid and 10 nmol internal standard are summarized in Supplementary Table 6c. The highest peak area ratio of free fatty acid to the internal standard (FFA/IS) was obtained for arachidonic acid (0.098). This in accordance with the ratio observed in the standard curve (Suppl. Fig. 10A). A lower FFA/IS was obtained for palmitic acid (0.026). As palmitic acid was not externally added to the control standard sample, it is assumed that palmitic fatty acid is ubiquitous present as a contamination in the laboratory materials. An FFA/IS value of 0.027 was obtained for a fatty acid, which co-elutes with nonadecanoic acid (C19:0). As this fatty acid was not externally added to the control standard sample nor it is expected to be a laboratory contamination, it can be hypothesized that it is a contamination in the commercially available preparation of the internal standard which is quasi a C19:0 fatty acid. The FFA/IS values of the other free fatty acids are remarkably lower (<0.0065), which suggest that they cannot be considered as appreciable contaminations (Suppl. Fig. 10B).

### Transmission Electron Microscopy

After dissection, brains of 15-day-old flies were fixed overnight immediately by immersion in 150 mM HEPES containing 1.5% glutaraldehyde and 1.5% formaldehyde at pH 7.35. Preparation for TEM was done as described (Mariani et al., 2022). Imaging was done in a Zeiss EM 900 at 80 kV, equipped with a side mount CCD camera (TRS).

## AKNOWLEDGMENTS

We would like to thank Doris Kretzschmar, Mikael Simons, Christian Klämbt, Hugo Bellen and Stanislava Chtarbanova-Rudloff for sharing flies and reagents with us. Marko Shcherbatyy for drawing a scheme. Christian Klämbt and Volkan Seyrantepe for contribution to phenotype description. All Shcherbata lab members for critical reading of the manuscript and helpful suggestions. The Medical School Hannover, VolkswagenStiftung (grants 90218 and 97750), and EMBO YIP for financial support.

## AUTHOR CONTRIBUTIONS

Conceptualization, HRS; Methodology, MIT, ASY, BB, DT, JH, HRS, Investigation, MIT, ASY, BB, DT, HRS, Writing, MIT, HRS, Visualization, MIT, ASY, HRS, Funding Acquisition, HRS.

## DECLARATION OF INTERESTS

The authors declare no competing interests.

This study includes no data deposited in external repositories.

## SUPPLEMENTARY INFORMATION

**Supplementary Figure 1.**
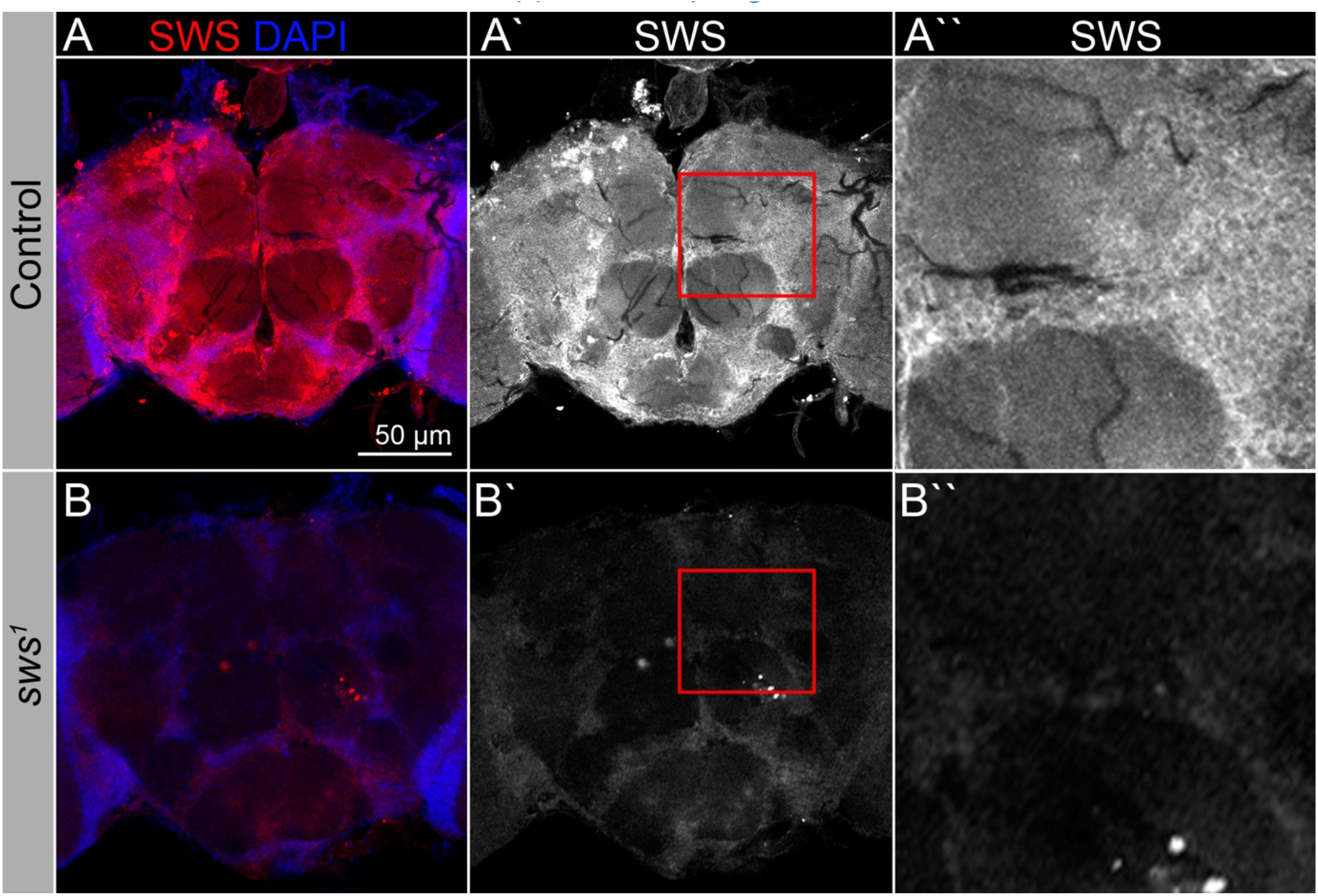
SWS expression pattern. **A-Aʹʹ.** Adult brains stained with anti-SWS antibodies (red) and DAPI (blue) show that SWS is expressed in most if not all brain cells in the control (*Oregon R*). **B-Bʹʹ.** In *sws^1^* mutant brains, SWS expression is dramatically reduced. Scale bar: 50 µm.

**Supplementary Figure 2.**
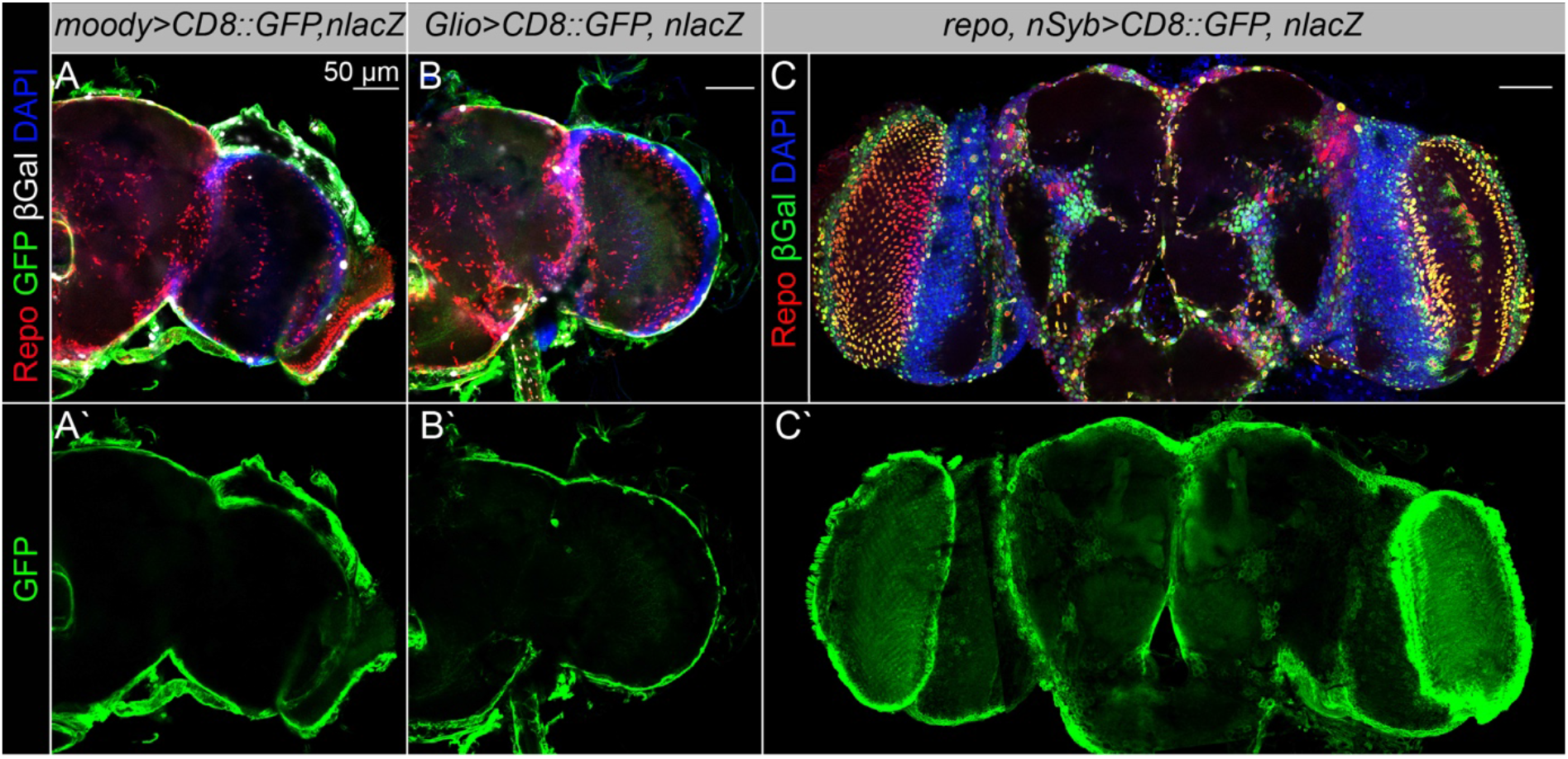
Expression patterns of the Gal4 driver lines used in the study. **A-Aʹ.** Expression pattern of *moody-Gal4* determined by combining of the transcriptional activator Gal4 under control of the *moody* gene promotor (*moody-Gal4*) and the *UAS-CD8::GFP* and *UAS-nLacZ* constructs. Fluorescence images of the brain show that *moody* is strongly expressed in the surface glia. Glia cells are marked with Repo (red), *moody* expression is indicated by the membrane CD8::GFP (green) and nuclear β-Galactosidase (βGal, white), and nuclei are marked with DAPI (blue). Scale bar: 50 µm. **B–Bʹ.** Expression pattern of *Gli-Gal4* determined by combining of the transcriptional activator Gal4 under control of the *Gliotactin* gene promotor (*Gli-Gal4*) and the *UAS-CD8::GFP* and *UAS-nLacZ* constructs. Fluorescence images of the brain show that *Gliotactin* is strongly expressed in the surface glia. Glia cells are marked with Repo (red), *Gli* expression is indicated by the membrane CD8::GFP (green) and nuclear β-Galactosidase (βGal, white), and nuclei are marked with DAPI (blue). Scale bar: 50 µm. **C–Cʹ.** Expression pattern of the double driver line (*repo, nSyb-Gal4*) determined by combining of the transcriptional activator Gal4 under control of the glial *repo* and neuronal *nSyb* promotors (*repo, nSyb-Gal4*) driving *UAS-CD8::GFP* and *UAS-nLacZ* transgenic constructs. Fluorescence images of the brain show that Repo and nSyb are strongly expressed thought the entire brain. Neuronal cell nuclei are marked by the nuclear β-Galactosidase expression driven by *nSyb-Gal4* (βGal, green), glial cell nuclei are marked by expression of the nuclear β-Galactosidase (βGal, green) driven by *repo-Gal4* and anti-Repo antibodies (red) (red+green=yellow) **(C)**. Fluorescence images of the brain show that *repo, nSyb-Gal4* is strongly expressed in glia and neurons marked by the membrane CD8::GFP (green) **(Cʹ)**. Scale bar: 50 µm.

**Supplementary Figure 3.**
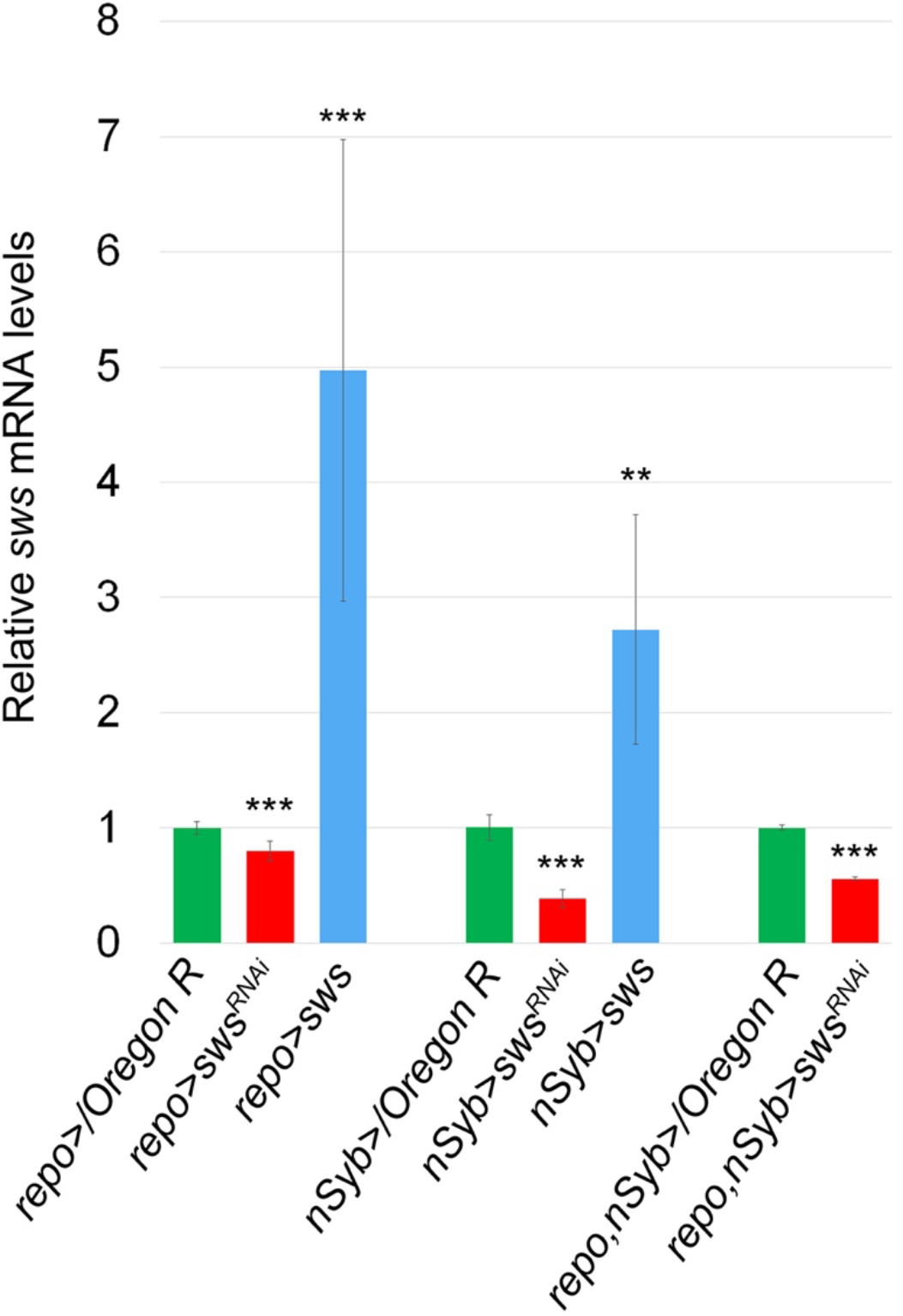
*sws* mRNA levels RT-qPCR analysis of *sws* mRNA levels from flies with glial, neuronal, or glial and neuronal *sws* downregulation (*repo>sws^RNAi^, nSyb>sws^RNAi^* and *repo, nSyb>sws^RNAi^*) confirms the efficacy of *sws^RNAi^* (red) and *UAS-sws* (blue) constructs. Two-tailed Student’s test was used to test for statistical significance - *repo>/OregonR* vs *repo>sws^RNAi^*, p=5.8E-4; *repo>/OregonR* vs *repo>sws,* p=6.7E-4; *nSyb>/Oregon R* vs *nSyb>sws^RNAi^*, p=5.2E-7; *nSyb>/OregonR* vs *nSyb>sws,* p=4.2E-3; *repo, nSyb>/OregonR vs repo, nSyb>sws^RNAi^*, p=7.9E-6.

**Supplementary Figure 4.**
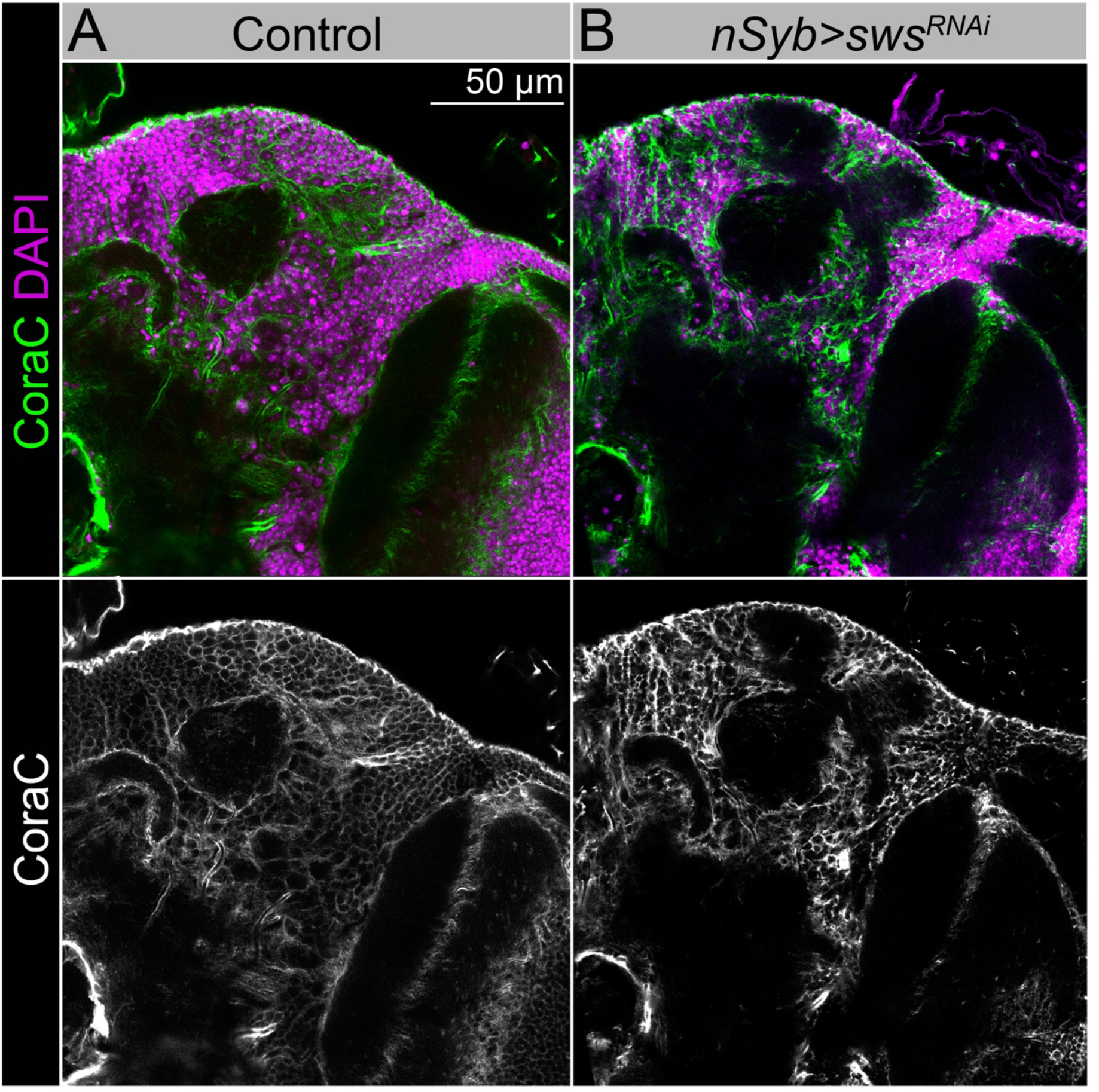
**A-B.** Adult brains stained with CoraC (green) and DAPI (magenta). Analysis of CoraC expression shows that, similar to control animals (**A,** *Oregon R x white^1118^*), animals with *sws* downregulation in neurons (**B,** *nSyb>sws^RNAi^*) do not have lesions and clumps in the brain surface. Scale bar: 50 µm.

**Supplementary Figure 5.**
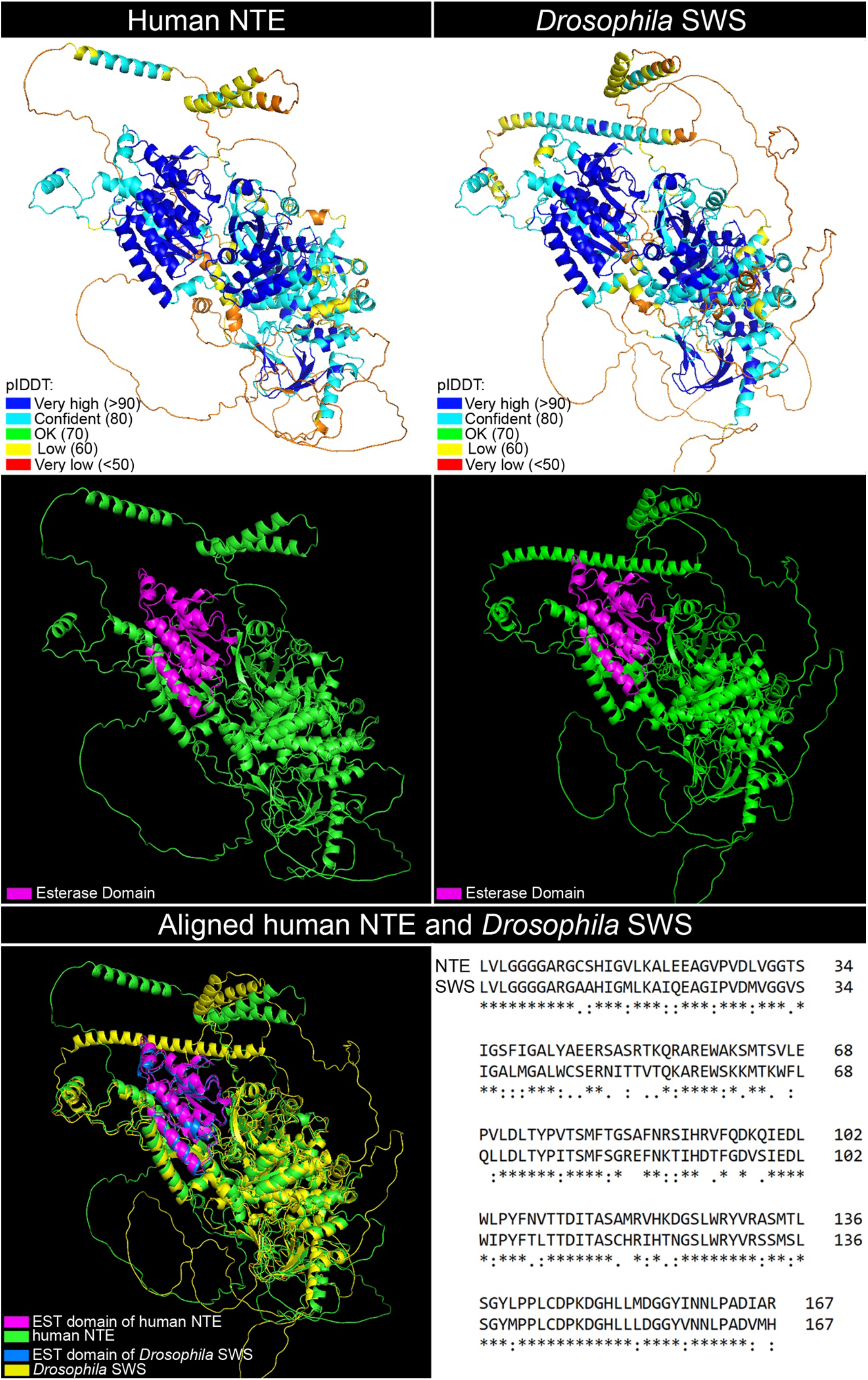
3D structures of human NTE and *Drosophila* SWS The 3D structures of the human NTE and Drosophila SWS proteins, generated using the AlphaFold2 and PyMOL tools. Both proteins contain a highly conserved patatin-like phospholipase domain known as the EST domain. In the SWS protein, this domain is located between amino acid residues 952 and 1118, while in the NTE protein, it spans residues 981 to 1147. The EST domain is characterized by a three-layer α/β/α architecture with a central six-stranded β-sheet sandwiched essentially between α-helices front and back. Comparison of the predicted structures of EST-SWS and EST-NTE show that they overlap. The EST domains in both proteins exhibit a high level of confidence as helices, with predicted local distance difference test scores (pLDDT) exceeding 90, indicating high accuracy and reliability in their structural predictions.

**Supplementary Figure 6.**
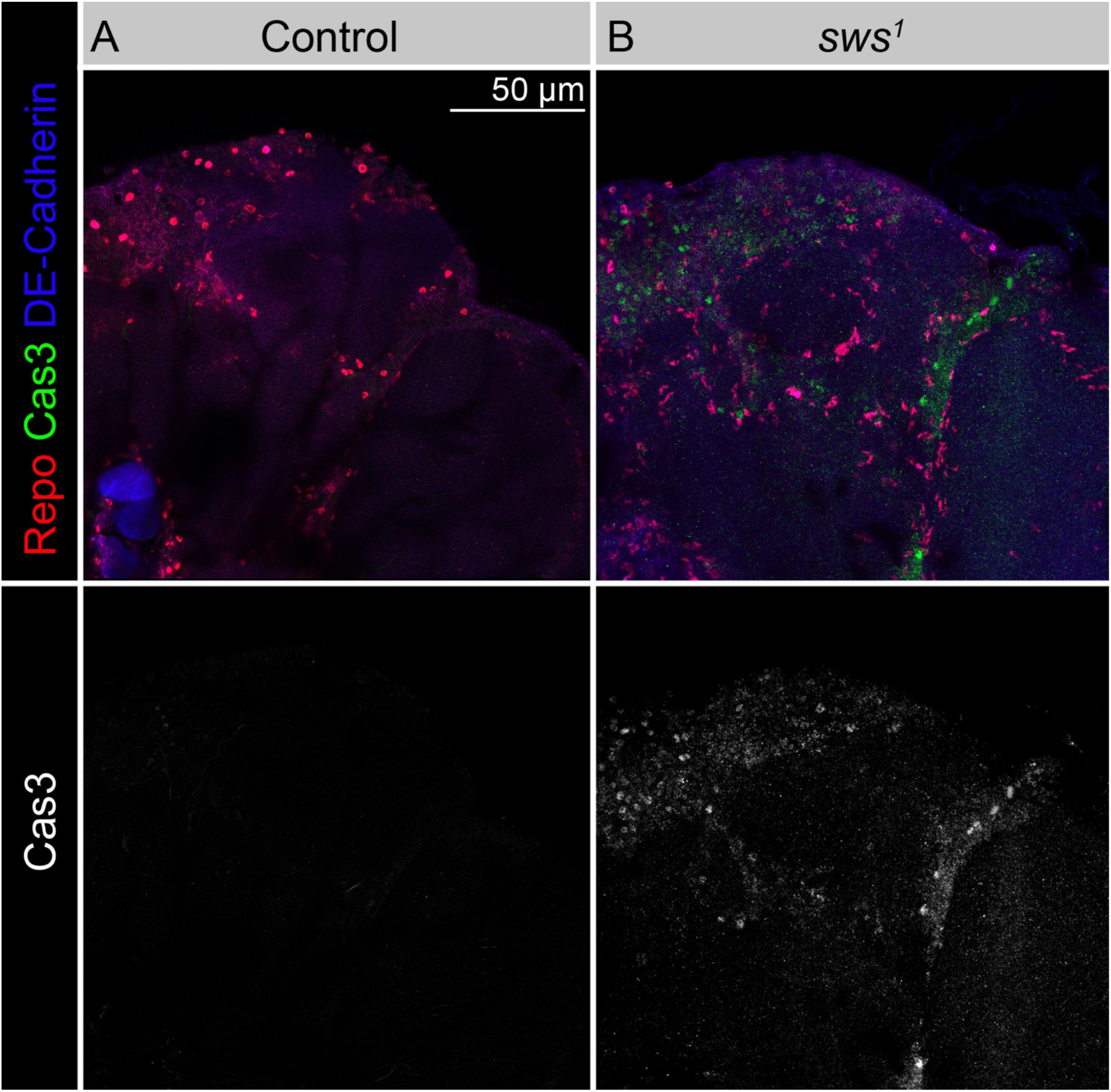
**A-B.** Adult brains stained with Repo (red, marks glia cell nuclei) and Cas3 (green, marks apoptotic cells) and DE-Cadherin (blue, marks cell membranes) to reveal apoptotic cell death in the brain. Note the increase in Cas3-positive cells (mainly neurons) inside *sws* mutant brains **(B)** when compared to *Oregon R* control **(A).**

**Supplementary Figure 7.**
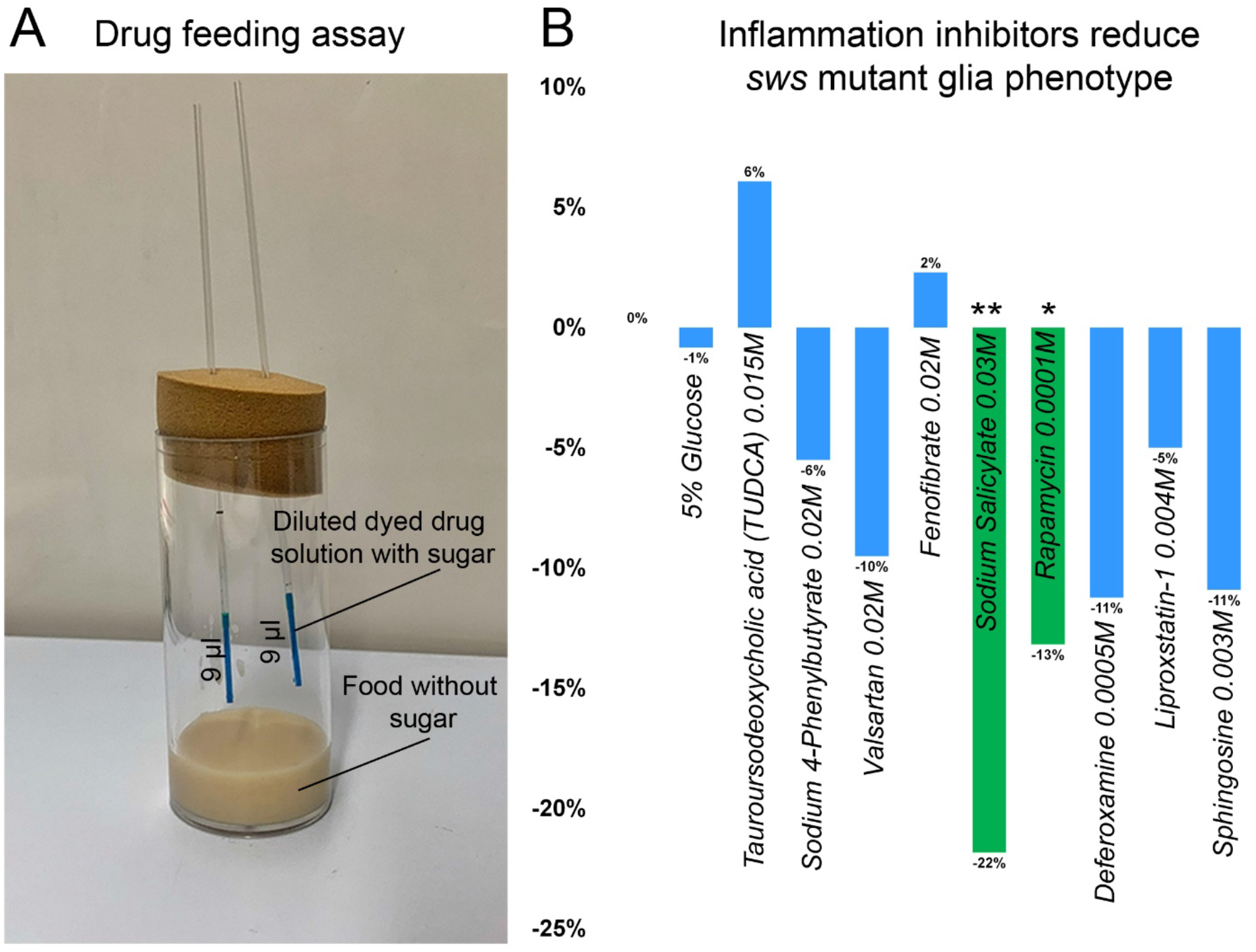
**A.** For the drug feeding assay, vials with sugar-free food with 2 micropipettes filled with dyed drug solution were used. **B.** Bar graph shows the changed percentage of the brains of *sws* mutants with the glial phenotype, assayed with CoraC, which were treated with different stress and inflammation inhibitors in comparison to untreated mutants. Two-way tables and chi-square test were used for statistical analysis. *p < 0.05, **p < 0.005, ***p < 0.001, n of adult brain hemispheres > 73, at least three biological replicates.

**Supplementary Figure 8.**
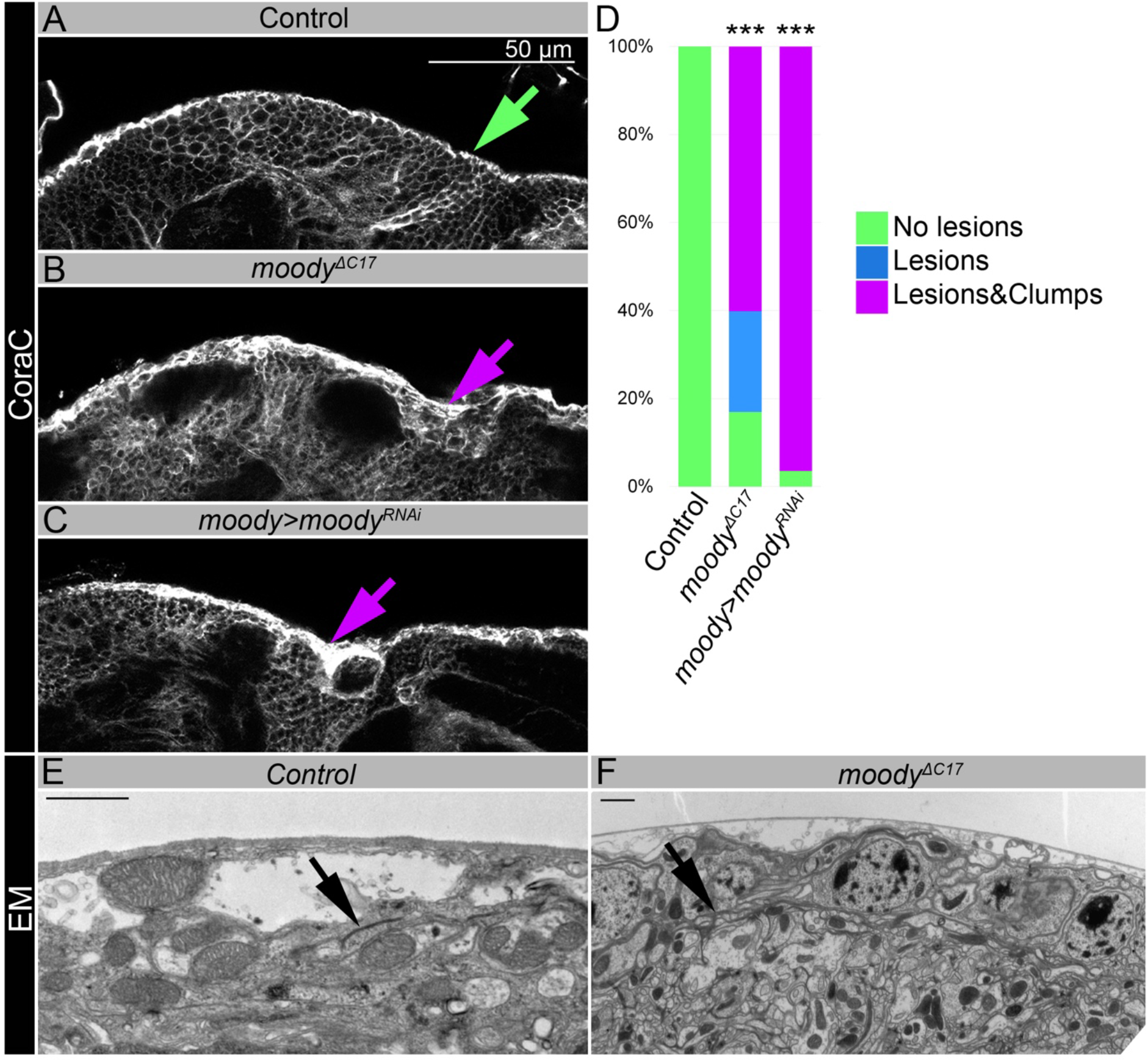
**A-C.** Adult brains stained with CoraC (white) to reveal brain surface. CoraC expression in control brains (*Oregon R x white^1118^*, **A)** is depicted as the smooth line at the surface of the brain (green arrow). In *moody^ΔC17^*mutant brains **(B)** and upon *moody* downregulation in SPG cells (*moody>moody^RNAi^*) **(C)** CoraC-positive outer cell layer contains lesions and clumps (magenta arrows). **A. D.** Bar graph shows the percentage of the brains with a defective brain surface. Two-way tables and chi-square test were used for statistical analysis, *** p < 0.001, n of adult brain hemispheres > 20. **E-F.** Electron microscope images of adult brain surface area of control (*white^1118^*) brain **A. (E)** and *moody^ΔC17^* mutant brain **(F)** depicted as the smooth line at the surface of the brain without any abnormal structures. Arrows show septate junctions in control **(E)** and *moody^ΔC17^* mutant brains **(F)**.

**Supplementary Figure 9.**
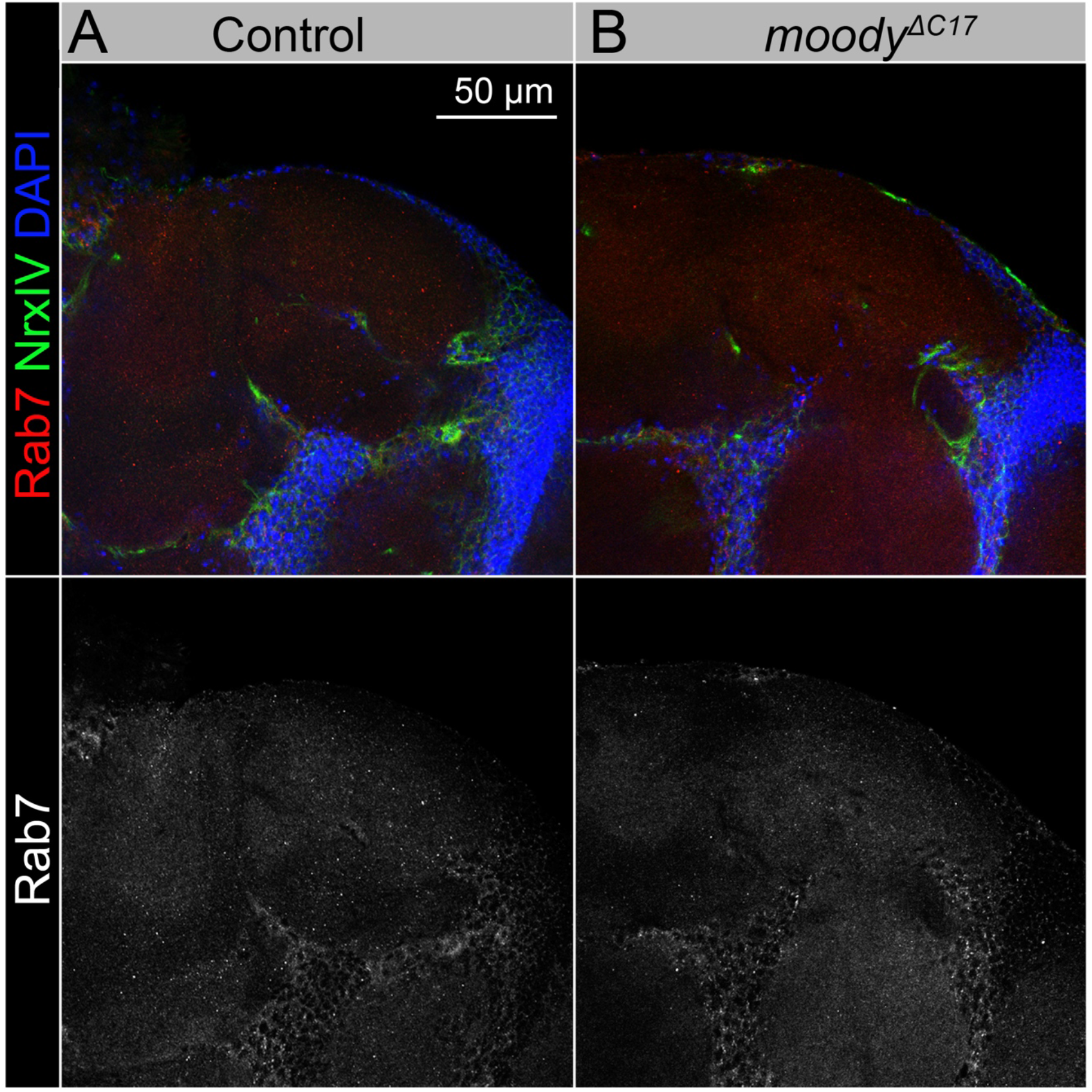
**A-B.** Adult brains stained with Rab7 (red) and NrxIV (green) and DAPI (blue) to reveal Rab7 vesicles in the brain, indicating that *Oregon R* control brain **(A)** and *moody^ΔC17^* mutant **(B)** did not show any abnormal accumulation of Rab7.

**Supplementary Figure 10.**
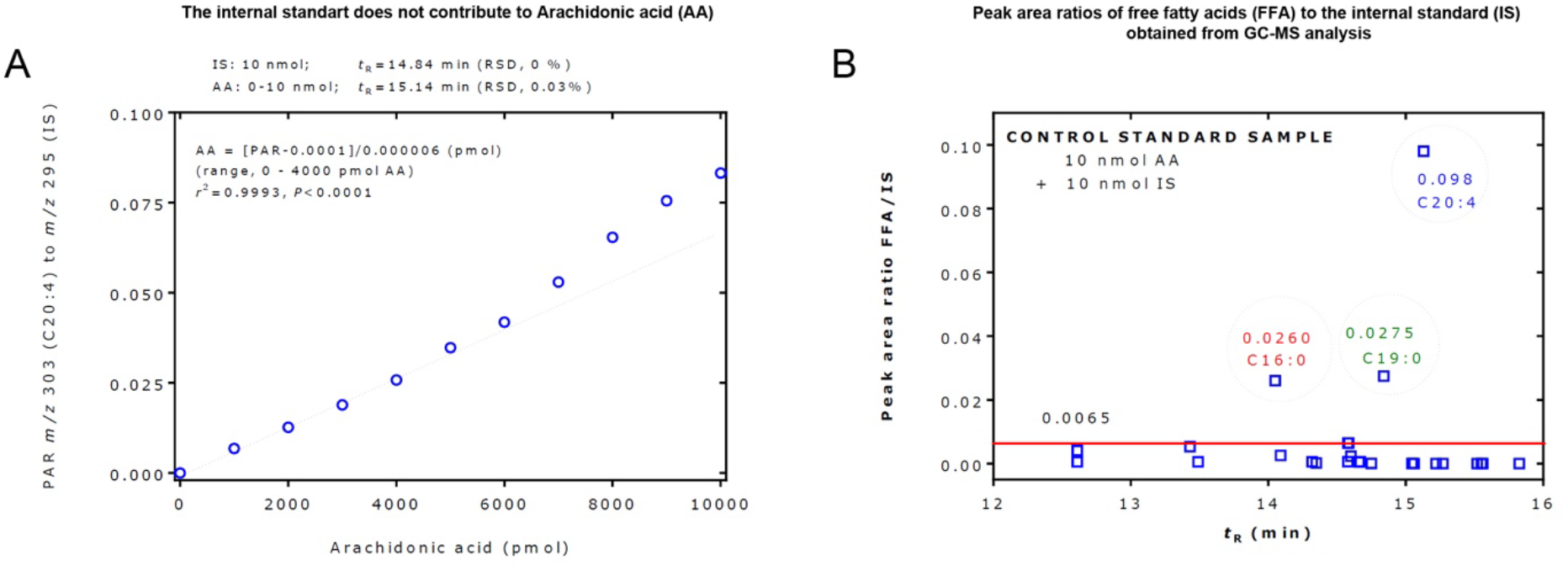
**A.** GC-MS measurement of arachidonic acid (C20:4) with the internal standard (IS). The peak area ratio (PAR) of *m/z* 303 for 20:4 to *m/z* 295 for the IS was linear in the range 0 to 4000 pmol of C20:4 at the fixed amount of 10000 pmol of the IS. The graph shows an example for the quantitative measurement of arachidonic acid (C20:4) with the internal standard (IS). The peak area ratio of m/z 303 for 20:4 to m/z 295 for the IS was linear in the range 0 to 4000 pmol C20:4 at the fixed amount of 10000 pmol of the IS. C20:4 and IS were baseline-separated by chromatography (14.84 min; RSD, 0% vs. 15.14 min; RSD, 0.03%) and entirely by mass spectrometry (m/z 295 vs m/z 303). The IS was found not to contribute to C20:4 by contaminating arachidonic acids or by its 13C isotope. On a molar basis, C20:4 produced about 10 times lower peak areas than the IS in a relevant concentration range of arachidonic acid. **B.** Peak area ratios of free fatty acids (FFA) to the internal standard (IS) obtained from GC-MS analysis of a control standard samples that contained 10 nmol arachidonic acid (C20:4) and 10 nmol of the IS. This Figure was constructed with the data of Supplementary Table 6c. Each symbol represents a free fatty acid. The horizontal red line at a FFA/IS value of 0.0065 suggests that FFA/IS values higher than 0.0065 can be considered to present in the control standard sample and/or as laboratory contaminations. For more details see the text. *t*R, retention time.

**SUPPLEMENTARY TABLE 1.** SWS expression in the surface glia is important for the integrity of *Drosophila* BBB.

| *Genotype* | Phenotypes |  |  | P-value | Number of brain hemi-spheres analyzed |
| --- | --- | --- | --- | --- | --- |
|  | No lesions | Lesions | Lesions + Clumps |  |  |
| **Control**<br>***OR x w<sup>1118</sup>*** | 100% | 0% | 0% |  | 96 |
| ***sws<sup>1</sup>*** | 14% | 11% | 75% | <sup>a</sup>p = 9.9E-31 | 85 |
| ***sws<sup>1</sup>/sws<sup>4</sup>*** | 13% | 28% | 59% | <sup>a</sup>p = 3.3E-29 | 69 |
| ***repo, nSyb>sws<sup>RNAi</sup>*** | 7% | 24% | 69% | <sup>a</sup>p = 1.5E-31 | 62 |
| ***repo>/Oregon R*** | 99% | 1% | 0% | <sup>a</sup>p = 0.27 | 80 |
| ***repo>sws<sup>RNAi</sup>*** | 1% | 13% | 86% | <sup>b</sup>p = 1.2E-31 | 70 |
| ***sws<sup>1</sup>; repo>sws*** | 100% | 0% | 0% | <sup>b</sup>p = 0.46<br><sup>c</sup>p = 2.3E-24 | 43 |
| ***sws<sup>1</sup>; repo>hNTE*** | 100% | 0% | 0% | <sup>b</sup>p = 1<br><sup>c</sup>p = 1.6E-28 | 62 |
| ***moody>/Oregon R*** | 100% | 0% | 0% | <sup>a</sup>p = 1 | 67 |
| ***moody>sws<sup>RNAi</sup>*** | 15% | 15% | 70% | <sup>b</sup>p = 1.1E-22 | 72 |
| ***sws<sup>1</sup>; moody>sws*** | 98% | 2% | 0% | <sup>b</sup>p = 0.3<br><sup>c</sup>p = 4.3E-21 | 63 |
| ***sws<sup>1</sup>; moody>hNTE*** | 100% | 0% | 0% | <sup>b</sup>p = 1<br><sup>c</sup>p = 1.9E-21 | 61 |
| ***Gli>/Oregon R*** | 100% | 0% | 0% | <sup>a</sup>p = 1 | 58 |
| ***Gli>sws<sup>RNAi</sup>*** | 8% | 12% | 80% | <sup>b</sup>p = 2.4E-22 | 60 |
| ***sws<sup>1</sup>; Gli>sws*** | 98% | 0% | 2% | <sup>b</sup>p = 0.34<br><sup>c</sup>p = 5.5E-23 | 65 |
| ***sws<sup>1</sup>; Gli>hNTE*** | 100% | 0% | 0% | <sup>b</sup>p = 1<br><sup>c</sup>p = 8.9E-23 | 60 |

<sup>a</sup> – compared to control (*OR x w<sup>1118</sup>*)
<sup>b</sup> – compared to *Gal4-driver x OR*
<sup>c</sup> – compared to *Gal4-driver x UAS-sws<sup>RNAi</sup>*
The values are reported from experiments done in triplicates.
For statistical analyses of the observed phenotypes, two-way tables and $\chi^2$ -test were used.

**SUPPLEMENTARY TABLE 2a.** SWS deficit in the surface glia results in the accumulation of Rab7-positive structures.

| **Genotype** | **Rab7 accumulation in the surface glia** |  | **P-value** | **Number of brain hemispheres analyzed** |
| --- | --- | --- | --- | --- |
|  | **No** | **Yes** |  |  |
| ***moody>/Oregon R 1d*** | 96% | 4% |  | 71 |
| ***moody>sws<sup>RNAi</sup> 1d*** | 58% | 42% | <sup>a</sup>p = 9.3E-8 | 77 |
| ***moody>/Oregon R 15d*** | 95% | 5% | <sup>b</sup>p = 0.82 | 59 |
| ***sws<sup>1</sup>; moody>CD8::GFP 15d*** | 18% | 82% | <sup>a</sup>p = 2E-15 | 44 |
| ***moody>sws<sup>RNAi</sup> 15d*** | 31% | 69% | <sup>a</sup>p = 3.4E-13<br><sup>b</sup>p = 1E-3 | 62 |

<sup>a</sup> – compared to *Gal4-driver x OR* animals of the same age
<sup>b</sup> – compared to 1-day old animals of the same genotype
The values are reported from experiments done in triplicates.
For statistical analyses of the observed phenotypes, two-way tables and $\chi^2$ -test were used.

**SUPPLEMENTARY TABLE 2b.** SWS deficit in the surface glia result in the accumulation of CathepsinL-positive structures.

| **Genotype** | **CathepsinL accumulation in the surface glia** |  | **P-value** | **Number of brain hemispheres analyzed** |
| --- | --- | --- | --- | --- |
|  | **No** | **Yes** |  |  |
| ***moody>/Oregon R 1d*** | 90% | 10% |  | 80 |
| ***moody>sws<sup>RNAi</sup> 1d*** | 59% | 41% | <sup>a</sup>p = 7.6E-6 | 81 |
| ***moody>/Oregon R 15d*** | 89% | 11% | <sup>b</sup>p = 0.8 | 62 |
| ***sws<sup>1</sup>; moody>CD8::GFP 15d*** | 43% | 57% | <sup>a</sup>p = 2.3E-7 | 51 |
| ***moody>sws<sup>RNAi</sup> 15d*** | 37% | 63% | <sup>a</sup>p = 1E-8<br><sup>b</sup>p = 0.013 | 49 |

<sup>a</sup> – compared to *Gal4-driver x OR* animals of the same age
<sup>b</sup> – compared to 1-day old animals of the same genotype
The values are reported from experiments done in triplicates.
For statistical analyses of the observed phenotypes, two-way tables and $\chi^2$ -test were used.

**SUPPLEMENTARY TABLE 3.** SWS deficit in the surface glia results in permeable brain barrier.

| **Genotype** | **BBB permeability** |  | **P-value** | **Number of brain hemispheres analyzed** |
| --- | --- | --- | --- | --- |
|  | **Normal** | **Permeable** |  |  |
| **Control (*Or x w<sup>1118</sup>*)** | 75% | 25% |  | 96 |
| ***sws<sup>1</sup>*** | 12% | 88% | <sup>a</sup>p = 1.4E-18 | 94 |
| ***sws<sup>1</sup>/sws<sup>4</sup>*** | 2% | 98% | <sup>a</sup>p = 7.7E-18 | 54 |
| ***repo, nSyb>sws<sup>RNAi</sup>*** | 12% | 88% | <sup>a</sup>p=1.01E-13 | 56 |
| ***repo>/Oregon R*** | 66% | 34% | <sup>a</sup>p = 0.23 | 59 |
| ***repo>sws<sup>RNAi</sup>*** | 9% | 91% | <sup>a</sup>p = 1.8E-14<br><sup>b</sup>p = 8.7E-10 | 53 |
| ***sws<sup>1</sup>; repo>sws*** | 18% | 82% | <sup>b</sup>p = 4E-4<br><sup>c</sup>p = 0.21 | 17 |
| ***sws<sup>1</sup>; repo>hNTE*** | 36% | 64% | <sup>b</sup>p = 2.8E-3<br><sup>c</sup>p = 1.4E-3 | 44 |
| ***moody>/Oregon R*** | 66% | 34% | <sup>a</sup>p = 0.17 | 91 |
| ***moody>sws<sup>RNAi</sup>*** | 9% | 91% | <sup>a</sup>p = 4.1E-16<br><sup>b</sup>p = 2.4E-12 | 64 |
| ***sws<sup>1</sup>; moody>sws*** | 22% | 78% | <sup>b</sup>p = 1.9E-8<br><sup>c</sup>p = 0.046 | 73 |
| ***sws<sup>1</sup>; moody>hNTE*** | 16% | 84% | <sup>b</sup>p = 1.4E-8<br><sup>c</sup>p = 0.29 | 50 |
| ***Gli>/Oregon R*** | 65% | 35% | <sup>a</sup>p = 0.15 | 71 |
| ***Gli>sws<sup>RNAi</sup>*** | 8% | 92% | <sup>a</sup>p = 2.1E-16<br><sup>b</sup>p = 1.9E-11 | 62 |
| ***sws<sup>1</sup>; Gli>sws*** | 25% | 75% | <sup>b</sup>p = 3.9E-6<br><sup>c</sup>p = 0.01 | 61 |
| ***sws<sup>1</sup>; Gli>hNTE*** | 26% | 74% | <sup>b</sup>p = 2.6E-5<br><sup>c</sup>p = 0.01 | 50 |

<sup>a</sup> – compared to control (*OR x w<sup>1118</sup>*)
<sup>b</sup> – compared to *Gal4-driver x OR*
<sup>c</sup> – compared to *Gal4-driver x UAS-sws<sup>RNAi</sup>*
The values are reported from experiments done in triplicates.
For statistical analyses of the observed phenotypes, two-way tables and $\chi^2$ -test were used.

**SUPPLEMENTARY TABLE 4.** moody^ΔC17^ flies show CoraC glial phenotypes similar to *sws* mutants.

| **Genotype** | **Phenotypes** |  |  | **P-value** | **Number of analyzed brain hemispheres** |
| --- | --- | --- | --- | --- | --- |
|  | **No lesions** | **Lesions** | **Lesions + Clumps** |  |  |
| **Control** | 100% | 0% | 0% |  | 96 |
| ***moody*<sup>ΔC17</sup>** | 17% | 23% | 60% | <sup>a</sup>p = 1.2E-32 | 118 |
| ***moody*>*moody*<sup>RNAi</sup>** | 4% | 0% | 96% | <sup>a</sup>p = 1.5E-27 | 28 |

<sup>a</sup> – compared to control (*OR* x *w*<sup>1118</sup>)
The values are reported from experiments done in triplicates.
For statistical analyses of the observed phenotypes, two-way tables and $\chi^2$ -test were used.

**SUPPLEMENTARY TABLE 5.**
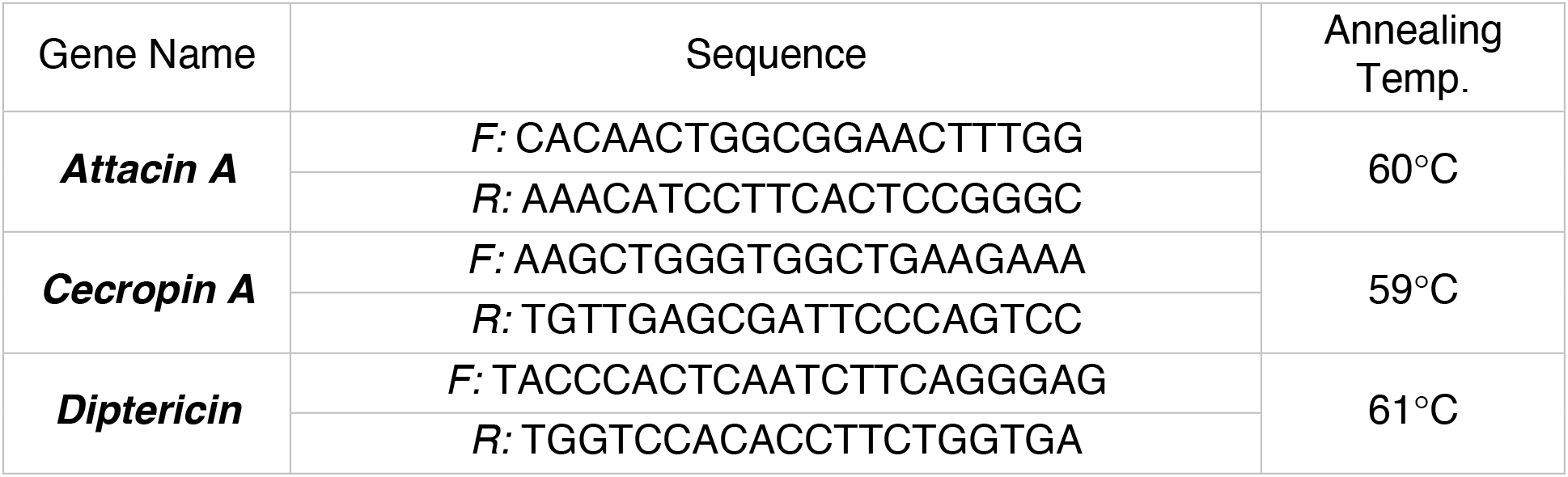
The list of AMP primers.

**SUPPLEMENTARY TABLE 6a.** sws and *moody* mutants show upregulated levels of free fatty acids.

| **Genotype** | **C14:1** | **C16:0** | **C16:1** | **C18:0** | **C18:1** | **C18:2** | **C18:3** | **C20:0** | **C20:4** | **C20:5** |
| --- | --- | --- | --- | --- | --- | --- | --- | --- | --- | --- |
|  | *FA/IS by weight (Average)* |  |  |  |  |  |  |  |  |  |
| **Control Oregon R** | 1.7x10<sup>2</sup> | 8.5x10<sup>2</sup> | 1.1x10<sup>3</sup> | 3.2x10<sup>2</sup> | 7.8x10<sup>2</sup> | 7.2x10<sup>2</sup> | 78.0x10<sup>2</sup> | 1.9 | 6.3x10<sup>-3</sup> | 6.4x10<sup>-3</sup> |
| ***sws*<sup>1</sup>** | 3.1x10<sup>2</sup> | 2.4x10<sup>3</sup> | 2.7x10<sup>3</sup> | 6.3x10<sup>2</sup> | 1.9x10<sup>3</sup> | 1.8x10<sup>3</sup> | 1.4x10<sup>3</sup> | 6.8 | 1.7x10<sup>-2</sup> | 9.4x10<sup>-3</sup> |
| **P-value** | 1E-6 |  |  |  |  |  |  |  |  |  |
| **Genotype** | **C14:1** | **C16:0** | **C16:1** | **C18:0** | **C18:1** | **C18:2** | **C18:3** | **C20:0** | **C20:4** | **C20:5** |
|  | *FA/IS by weight (Average)* |  |  |  |  |  |  |  |  |  |
| **Control white<sup>1118</sup>** | 2.3x10<sup>2</sup> | 1.1x10<sup>3</sup> | 1.5x10<sup>3</sup> | 3.7x10<sup>2</sup> | 1.1x10<sup>3</sup> | 9.5x10<sup>2</sup> | 1.1x10<sup>3</sup> | 2.1 | 8.6x10<sup>-3</sup> | 5.8x10<sup>-3</sup> |
| ***moody* ΔC17** | 2.7x10<sup>2</sup> | 2.0x10<sup>3</sup> | 2.9x10<sup>3</sup> | 6.1x10<sup>2</sup> | 2.2x10<sup>3</sup> | 2.0x10<sup>3</sup> | 2.2x10<sup>3</sup> | 4.1 | 2.0x10<sup>-2</sup> | 1.3x10<sup>-2</sup> |
| **P-value** | 6.7E-9 |  |  |  |  |  |  |  |  |  |

For statistical analyses one-way ANOVA test was used.
C14:1 - 9-cis-Tetradecenoic acid
C16:0 - Palmitic acid
C16:1 - Palmitoleic acid
C18:0 - Stearic acid
C18:1 - Oleic acid
C18:2 - Linoleic acid
C18:3 - $\alpha$ - and $\gamma$ -Linolenic acid
C20:0 - Eicosanoic acid

**SUPPLEMENTARY TABLE 6b.** Free fatty acids analyzed by GC-MS after derivatization to their pentafluorobenzyl esters.

| Fatty acid | Acronym | MW<br>(Da) | t<sub>R</sub><br>(min) | SIM<br>(m/z) |
| --- | --- | --- | --- | --- |
| Dodecanoic acid (lauric acid) | C12:0 | 200.3 | 12.62 | 199 |
| (5Z)-Dodecenoic acid | C12:1 | 198.3 | 12.65 | 197 |
| Tetradecanoic acid (myristic acid) | C14:0 | 228.4 | 13.44 | 227 |
| 9-cis-Tetradecenoic acid | C14:1 | 226.4 | 13.52 | 225 |
| Hexadecanoic acid (palmitic acid) | C16:0 | 256.4 | 14.07 | 255 |
| 9-cis-Hexadecenoic acid | C16:1 | 254.4 | 14.10 | 253 |
| Heptadecanoic acid | C17:0 | 270.4 | 14.31 | 269 |
| Heptadecenoic acid | 10c-C17:1 | 268.4 | 14.39 | 267 |
| Octadecanoic acid (stearic acid) | C18:0 | 284.5 | 14.58 | 283 |
| 9-cis-Octadecenoic acid (oleic acid) | C18:1 | 282.5 | 14.62 | 281 |
| 9-trans-Octadecenoic acid | 9t-C18:1 | 282.5 | 14.62 | 281 |
| 11-trans-Octadecenoic acid | 11t-C18:1 | 282.5 | 14.64 | 281 |
| 6-cis-Octadecenoic acid | 6c-C18:1 | 282.5 | 14.60 | 281 |
| Linoleic acid | C18:2 | 280.5 | 14.68 | 279 |
| alpha-Linolenic acid | C18:3 | 278.4 | 14.76 | 277 |
| gamma-Linolenic acid | C18:3 | 278.4 | 14.68 | 277 |
| Nonadecanoic acid | C19:0 | 298.5 | 14.83 | 297 |
| **INTERNAL STANDARD** | **8-cp-C18:0** | **296.5** | **14.81** | **295** |
| Eicosanoic acid | C20:0 | 312.5 | 15.05 | 311 |
| 11-cis-Eicosenoic acid | 11c-C20:1 | 310.5 | 15.08 | 309 |
| Arachidonic acid (AA) | C20:4 | 304.5 | 15.13 | 303 |
| Eicosapentanoic acid (EPA) | C20:5 | 302.5 | 15.22 | 301 |
| Heneisosanoic acid | C21:0 | 326.6 | 15.26 | 325 |
| Docosanoic acid | C22:0 | 340.6 | 15.46 | 339 |
| Erucic acid | 13c-C22:1 | 338.6 | 15.52 | 337 |
| Tetracosanoic acid | C24:0 | 368.6 | 15.85 | 367 |
| 15-cis-Tetracosenoic acid | 15c-C24:1 | 366.6 | 15.99 | 365 |

**SUPPLEMENTARY TABLE 6c.** Mean peak area ratio and its coefficient of variation (CV) of the listed free fatty acids (FFA) to the internal standard (IS) in the diluted (1:10, v/v) control standard sample (10 nmol arachidonic acid, 10 nmol internal standard) analyzed in duplicate as pentafluorobenzyl esters.

| Fatty acid | Acronym | FFA/IS | CV (%) |
| --- | --- | --- | --- |
| Dodecanoic acid (lauric acid) | C12:0 | 4.3 E-03 | 13.8 |
| (5Z)-Dodecenoic acid | C12:1 | 6.0 E-04 | 15.1 |
| Tetradecanoic acid (myristic acid) | C14:0 | 5.3 E-03 | 10.7 |
| 9-cis-Tetradecenoic acid | C14:1 | 6.0 E-04 | 1.3 |
| Hexadecanoic acid (palmitic acid) | C16:0 | **2.6 E-02** | 7.7 |
| 9-cis-Hexadecenoic acid | C16:1 | 2.6 E-03 | 1.0 |
| Heptadecanoic acid | C17:0 | 6.0 E-04 | 1.3 |
| Heptadecenoic acid | 10c-C17:1 | 2.1 E-04 | 3.8 |
| Octadecanoic acid (stearic acid) | C18:0 | 6.6 E-03 | 11.1 |
| 9-cis-Octadecenoic acid (oleic acid) | C18:1 | 2.4 E-03 | 4.4 |
| 9-trans-Octadecenoic acid | 9t-C18:1 | not found |  |
| 11-trans-Octadecenoic acid | 1t-C18:1 | 2.3 E-03 | 2.0 |
| 6-cis-Octadecenoic acid | 6c-C18:1 | 2.3 E-03 | 2.0 |
| Linoleic acid | C18:2 | 6.1 E-04 | 2.7 |
| alpha-Linolenic acid | C18:3 | 5.7 E-05 | 16.4 |
| gamma-Linolenic acid | C18:3 | 5.6 E-05 | 15.7 |
| Nonadecanoic acid | C19:0 | **2.7 E-02** | 1.1 |
| **INTERNAL STANDARD** | **8-cp-C18:0** | **1:1** | **n.a.** |
| Eicosanoic acid | C20:0 | 6.1 E-05 | 16.0 |
| 11-cis-Eicosenoic acid | 11c-C20:1 | 5.6 E-05 | 38.7 |
| Arachidonic acid (AA) | C20:4 | **9.8 E-02** | 0.8 |
| Eicosapentanoic acid (EPA) | C20:5 | 5.2 E-05 | 21.5 |
| Heneisosanoic acid | C21:0 | 2.0 E-05 | 24.8 |
| Docosanoic acid | C22:0 | 1.9 E-05 | 13.1 |
| Erucic acid | 13c-C22:1 | 3.6 E-05 | 3.6 |
| Tetracosanoic acid | C24:0 | 4.9 E-06 | 27.6 |
| 15-cis-Tetracisenoic acid | 15c-C24:1 | not found |  |

**SUPPLEMENTARY TABLE 7.** The effect of treatment with different anti-inflammatory substances and stress suppressors on the frequency of the surface glia phenotype in *sws* and *moody* mutants.

| **Genotype + Drug** | **% of brain hemispheres with CoraC phenotype** | **Delta % after treatment** | **P-value** | **Number of brain hemispheres analyzed** |
| --- | --- | --- | --- | --- |
| ***sws*<sup>1</sup>** | 75% |  |  | 85 |
| ***sws*<sup>1</sup> + 5% Glucose** | 74% | -1% | <sup>a</sup>p = 0.9 | 94 |
| ***sws*<sup>1</sup> + Tauroursodeoxycholic acid (TUDCA) 0.015M** | 81% | 6% | <sup>a</sup>p = 0.31 | 102 |
| ***sws*<sup>1</sup> + Sodium 4-Phenylbutyrate 0.02M** | 70% | -6% | <sup>a</sup>p = 0.37 | 139 |
| ***sws*<sup>1</sup> + Valsartan 0.02M** | 66% | -10% | <sup>a</sup>p = 0.15 | 114 |
| ***sws*<sup>1</sup> + Fenofibrate 0.02M** | 78% | 2% | <sup>a</sup>p = 0.71 | 116 |
| ***sws*<sup>1</sup> + Sodium Salicylate 0.03M** | 53% | -22% | <sup>a</sup>p = 1E-3 | 144 |
| ***sws*<sup>1</sup> + Rapamycin 0.0001M** | 62% | -13% | <sup>a</sup>p = 0.05 | 132 |
| ***sws*<sup>1</sup> + Deferoxamine mesylate salt 0.0005M** | 64% | -11% | <sup>a</sup>p = 0.08 | 128 |
| ***sws*<sup>1</sup> + Liproxstatin-1 0.004M** | 70% | -5% | <sup>a</sup>p = 0.45 | 101 |
| ***sws*<sup>1</sup> + Sphingosine 0.003M** | 64% | -11% | <sup>a</sup>p = 0.14 | 73 |
| ***sws*<sup>1</sup> + 5%Glucose** | 79% | 0% |  | 134 |
| ***sws*<sup>1</sup> + Sodium Sal. 0.03M** | 49% | -30% | <sup>a</sup>p = 4E-7 | 130 |
| ***sws*<sup>1</sup> + Rapamycin 0.0001M** | 62% | -17% | <sup>a</sup>p = 1.8E-3 | 133 |
| ***moody*<sup>ΔC17</sup> + 5%Glucose** | 60% | 0% |  | 118 |
| ***moody*<sup>ΔC17</sup> + Sodium Sal. 0.03M** | 46% | -14% | <sup>b</sup>p = 0.02 | 156 |
| ***moody*<sup>ΔC17</sup> + Rapamycin 0.0001M** | 39% | -21% | <sup>b</sup>p = 2E-3 | 104 |

<sup>a</sup> – compared to *sws*<sup>1</sup> (no drug treatment)
<sup>b</sup> – compared to *moody*<sup>ΔC17</sup> (no drug treatment)
The values are reported from experiments done in triplicates.
For statistical analyses of the observed phenotypes, two-way tables and $\chi^2$ -test were used.

## Notes

### Competing Interest Statement

The authors have declared no competing interest.

